# Deep mutational scanning of an *S. pneumoniae* FMN riboswitch reveals diverging adaptive landscapes in response to targeting antibiotics and robustness during infection

**DOI:** 10.1101/2025.04.17.649428

**Authors:** Rebecca Korn, Laurie Brutus, Jon S. Anthony, Quinlan Furumo, Elizabeth C. Gray, Indu Warrier, Michelle M. Meyer

## Abstract

RNA-based control of gene expression is common across all domains of life, yet shows extraordinary variety in terms of environmental cues recognized, mechanisms of action, required factors, and magnitude of expression changes. The fitness benefit conferred by RNA-mediated regulation is often subtle and challenging to measure and may be highly influenced by the environment. Yet selective forces are sufficient to generate and maintain complex RNA structures that allow rapid and robust response to stimuli. In this work we perform deep mutational scanning to measure the relative fitness conferred by all single mutants of an FMN riboswitch to the opportunistic pathogen *Streptococcus pneumoniae* under seven different conditions. We find the RNA sequence is not under selective pressure in standard planktonic culture. However, in the presence of two different targeting antibiotics diverging adaptive landscapes are exposed that reveal suites of mutations conferring either constitutive activation or repression of gene expression. Analysis of these landscapes highlights a complex relationship between antibiotic concentration and the fitness benefit conferred by individual mutations as well as illuminates the mechanism of action for this RNA. Finally, assessment within mouse lung infection reveals a sequence subject to widespread weak purifying selection, but robust to strongly deleterious mutations.

## INTRODUCTION

The interplay between genotype and phenotype is central to understanding how organisms adapt to their environments, evolve new traits, and respond to external challenges. In bacteria, regulation that occurs after transcription initiation plays an important role in allowing rapid adjustment to environmental changes. Regulatory RNAs such as trans-acting small RNAs (sRNAs) and cis-acting elements, including riboswitches and RNA thermometers, modulate gene expression in response to environmental cues, tuning metabolic processes, inducing stress responses, and activating resistance mechanisms^1^. Riboswitches, RNAs that respond to specific small molecule metabolites to alter gene expression, often control essential metabolic pathways and transporters. These RNAs act as molecular sensors that integrate metabolic cues with regulatory features to allow conditional intrinsic transcription termination or translation initiation. Riboswitches have emerged as potential targets for antimicrobials^1–8^, and due to their dynamic nature and relatively small size, have also proven to be valuable biophysical model systems for studying the relationships between RNA sequence, structure, and function *in vitro*^9,10^. However, our understanding of how sequence changes in such RNAs drive functional changes in regulation within the cell, and how these regulatory shifts ultimately affect an organism’s phenotype and fitness in diverse or dynamic environments, remains limited.

Fitness landscapes measured for other RNAs have yielded invaluable insights into the RNA folding landscape^11^, the effects of epistasis^12^, the degree to which fitness peaks are isolated from one another^13–15^, and the impacts of environmental perturbations on RNA function^16,17^. However, such studies are generally conducted *in vitro* to enable assessment of large numbers of variants^11,13–17^ or utilize engineered genetic selection schemes to partition functional and non-functional sequences^12^. Thus, how changes in ncRNA sequence influence regulatory function and ultimately organism phenotype remain unstudied in the context of dynamic regulatory RNAs that are dependent on cellular components for biological function.

In this work, we generated fitness landscapes for a 372 nucleotide *Streptococcus pneumoniae* flavin mononucleotide (FMN) riboswitch^18^ that assess all 1,116 single point mutations under seven different conditions including *in vivo* infection and antibiotic perturbed environments. *S. pneumoniae* is a significant global health concern as the leading cause of lower respiratory infection worldwide, contributing to over 1 million deaths globally in 2016^19^. In the United States, antibiotic-resistant pneumococcal diseases cause around 1.2 million illnesses and 7,000 deaths annually^20^. There are two FMN riboswitches in *S. pneumoniae* that function as cis-regulatory “off” switches; one preceding the riboflavin biosynthesis operon (*ribB/A,D,E,H*) and the other preceding the gene encoding the riboflavin transport protein RibU. Riboflavin (vitamin B_2_) is an essential nutrient, and its derivatives, FMN and flavin adenine dinucleotide (FAD), serve as critical cofactors in cellular metabolism, catalyzing redox reactions^1^. The FMN riboswitch is one of the most frequent and widely distributed riboswitches, occurring across a broad swath of diverse bacterial clades, with especially high prevalence across potential pathogenic species^21,22^. Thus, this riboswitch is specifically targeted by several different antimicrobial compounds that inhibit both FMN synthesis and transport to starve cells of flavin co-factors^23–29^.

We show that the FMN riboswitch preceding *ribU* is under minimal selection in standard planktonic culture conditions. However, antibiotic perturbation of culture conditions identified suites of mutations with constitutively active and repressed behavior, which in turn enable insights into the RNA mechanism of action and routes to drug tolerance. Furthermore, within *in vivo* mouse lung infections the RNA is under weak purifying selection, and *S. pneumoniae* fitness is robust to single mutations, even those with substantial impacts on gene expression. Our study uniquely assesses a regulatory RNA function within its native context to highlight the complex interplay between environment, selective pressure on the organism from the environment, and potential phenotypes accessible via single mutations.

## RESULTS

### Single mutations to the FMN riboswitch are neutral in planktonic culture in the presence and absence of riboflavin

The *S. pneumoniae* FMN riboswitch preceding *ribU* spans 372 nucleotides, from the transcription start site to a putative transcription termination site just upstream of the translational start identified via 3’-end sequencing^30^ **(Fig. 1A, Fig. S1).** The riboswitch includes a canonical six-stem FMN-binding aptamer with an extended P3 stem, followed by an extended stem after the aptamer that appears to enable intrinsic transcription termination, although the base-pairing changes required to connect ligand binding to termination are unclear **(Fig. 1A)**. To assess the impact of each of the 1,116 possible single-point mutants on *S. pneumoniae* fitness, we created a library of *S. pneumoniae* strains that integrates these mutations into the native locus accompanied by a chloramphenicol marker upstream of the native promoter sequence^31^. Our library maximizes the proportion of single point-mutations via combination of six individually constructed libraries, each of which contains a single segment comprised of a synthetic oligo doped with 1% of each alternative base at each position (see **Fig. S2A** for construction methodology).

**Figure 1:**
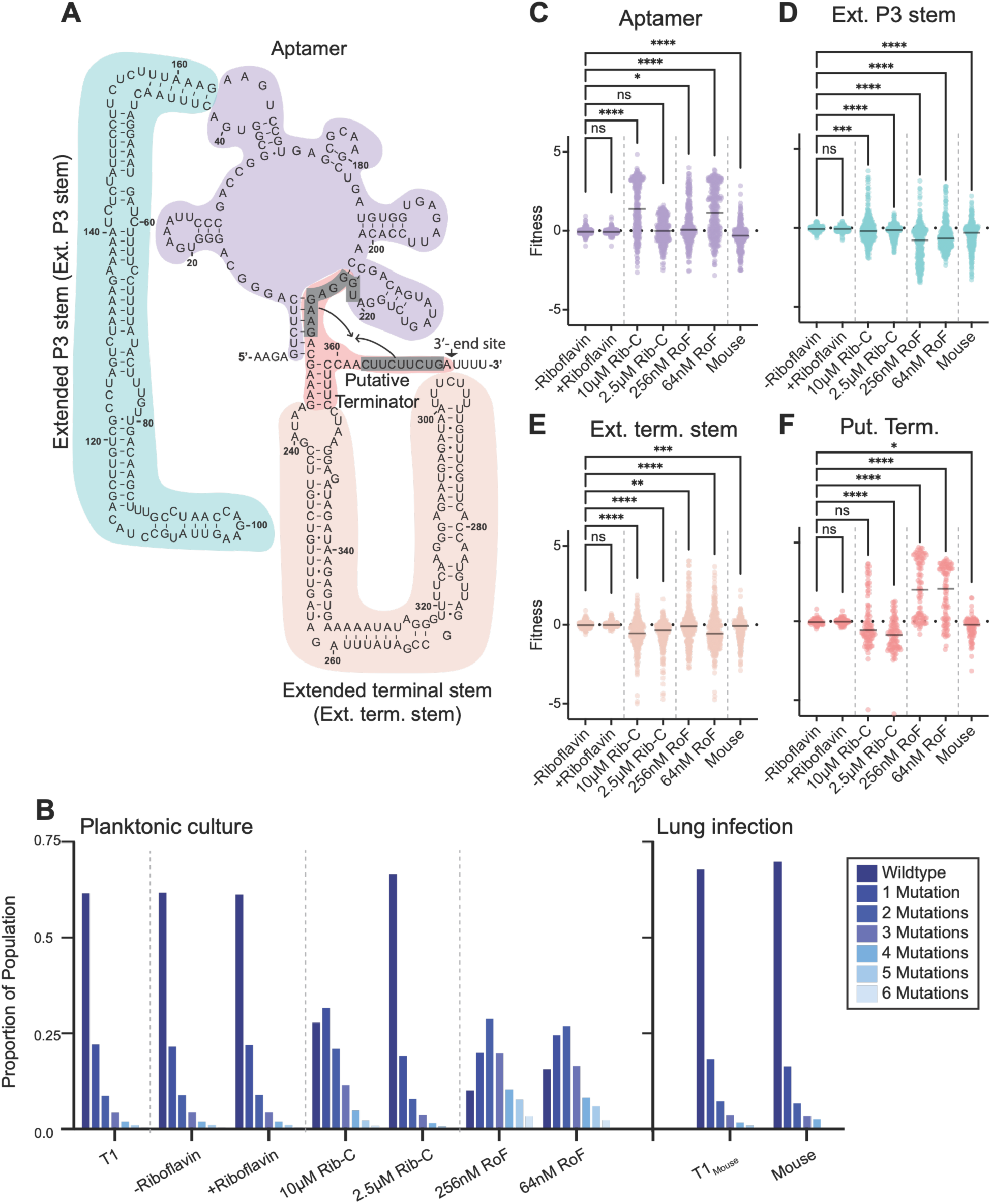
The mutant strain library is under selective pressure when challenged with antibiotics or within lung infection. **A)** The SP_0488 FMN riboswitch secondary structure colored according to region. Secondary structure derived from alignment to FMN aptamer consensus structure^43^, and folding extended stems with the RNAfold webserver^51^. 3’-end from previously collected Term-seq data^30^ is indicated (Fig. S1). **B)** The proportion of processed reads in each post-selection (T2) condition with 0 (wildtype sequence) to 6 mutations and their respective unselected (T1) distribution. **C-F)** Fitness of mutations to different regions of the riboswitch (color coded in A) relative to the wildtype sequence in seven different environments: chemically defined media (CDM) with no riboflavin (-Riboflavin), CDM+ 372 nM riboflavin (+Riboflavin), CDM+ 10µM ribocil-C (10µM Rib-C), CDM+ 2.5µM ribocil-C (2.5µM Rib-C), CDM+ 256nM roseoflavin (256nM RoF), CDM+ 64nM roseoflavin (64nM RoF), and within a mouse lung infection (Mouse). For each region, the fitness of the population of single mutants is compared to the fitness in the −Riboflavin condition using Kruskal-Wallis test with Dunn’s test for multiple comparisons (adj. p>0.05 (ns), adj. p<0.05 (*), adj. p<0.01 (**), adj. p<0.001 (***), p<0.0001 (****)). Population medians and exact p-values for each comparison on Table S3.

With this library we assessed the relative fitness of each strain compared to the wild-type sequence in a chemically defined medium in the presence and absence of riboflavin (CDM + and −riboflavin), which is rapidly converted into FMN via phosphorylation inside the cell. We cultured our strain library to mid-log phase and split the culture to harvest two pre-selection samples (T1 samples) and to inoculate cultures in CDM + and −riboflavin (each in technical triplicate) that were allowed to grow ∼10 doublings before harvest (T2 samples). Genomic DNA was extracted from both T1 and T2 samples, and the FMN riboswitch preceding *ribU* amplified via PCR with unique molecular identifiers (UMIs) to enable accurate quantification of each variant. The resulting amplicons were sequenced to obtain 0.6 - 7.3 million reads for each sample **(Fig. S2B, Fig. S3, Table S1)**. In both the presence and absence of riboflavin, the distribution of wildtype versus mutants with 1-6 mutations is similar across the T1 and T2 samples, with about 60% of reads corresponding to the wildtype sequence **(Fig. 1B)**, and all single point mutations are represented in the datasets at both the T1 and T2 conditions **(Fig. S2C, Table S2)**. The fitness of double mutants was not assessed because the number of double mutants is large (1,245,456), and our construction methodology does not create all pairs of double mutants.

To quantify the fitness of each single mutant, we used the DiMSum pipeline^32^ to analyze 1.0 – 12.5 million deconvoluted reads for each condition (**Fig. S3A-C, Table S2)**. DimSum calculates fitness relative to the wild-type sequence based on the frequency of each mutant relative to wild-type (see Methods), where a fitness of 0 is equivalent to the wild-type sequence, a fitness >0 is increased frequency in T2, and a fitness <0 is a decreased frequency in T2. We find that in both the presence and absence of riboflavin, no individual single-point mutation exhibited a fitness greater than 1 (beneficial) or less than −1 (detrimental) (**Fig. 1C-F, Table S2**), and the median fitness across all mutants is −0.052 and −0.045 in the absence and presence of riboflavin, respectively. We assessed the fitness of transversions (purine to pyrimidine mutation) and transitions (purine to purine or pyrimidine to pyrimidine mutation) to determine if changes more likely to be disruptive of Watson-Crick pairing (transversions) have a greater impact on fitness. Transversions show slightly lower fitness than transitions across all regions of the sequence, however, the median fitness difference is very small (Δmedian<0.1) (**Figure S4A-D**) suggesting that these differences are likely not biologically significant in the context of our experiment. Thus, consistent with minimal change in our distribution of mutation frequency, point mutations do not confer a substantial fitness advantage or disadvantage to *S. pneumoniae* in standard culture conditions **(Fig. 1B)**. Constitutive expression of the transporter at levels accessible by single mutations is not detrimental, and the distally located intact riboflavin biosynthetic operon (SP_0178 - SP_0175) provides sufficient riboflavin for any individuals displaying constitutive repression of RibU.

### Mutations disruptive to the FMN aptamer are beneficial in the presence of ribocil-C

Due to the prevalence of the FMN riboswitch across human bacterial pathogens, it has been the subject of drug development and range of diverse compounds have been identified that specifically interact with the FMN aptamer^24,25,28,33^. These compounds result in constitutive repression of genes in both FMN synthesis and transport to ultimately starve the cell of necessary flavin cofactors inhibiting growth. Such antibiotics include roseoflavin, a natural product structural analog of riboflavin, and ribocil-C, a synthetic molecule identified via high-throughput screening that also binds to the RNA competitively with FMN. Due to its inhibitory action on the biosynthetic operon, ribocil-C also enables us to probe how mutations to the *ribU* riboswitch impact fitness while FMN synthesis is inhibited.

To determine the full spectrum mutations to the FMN riboswitch preceding *ribU* that confer fitness changes in the presence of ribocil-C, the mutant strain library was grown in CDM −riboflavin supplemented with 2.5µM and 10µM ribocil-C similarly to the experiments above. The ribocil-C concentrations were chosen to be above (2.5µM), and substantially above (10µM), the IC50 of ribocil-C for wildtype *S. pneumoniae* TIGR4 (∼1µM). The growth of the strain library was inhibited in these conditions, with the cultures reaching OD_600_ of ∼0.8 and <0.4 respectively after 12.5 hours of growth (∼8-9 doublings). At 2.5 µM ribocil-C, the distribution of wild-type and mutated variants is similar to our T1 and non-selective conditions. However, at 10µM ribocil-C the distribution of wild-type and mutated variants was altered with the majority of reads containing 1 or more mutations **(Fig. 1B)**. Directly comparing the fitness values of all mutations at the two ribocil-C concentrations shows that they are highly correlated, but the magnitude of positive selection at 2.5µM ribocil-C is less than that of 10µM ribocil-C, consistent with the shift away from the wildtype sequence at higher antibiotic concentration (**Fig. 2A, Fig. S4E,F**)

**Figure 2:**
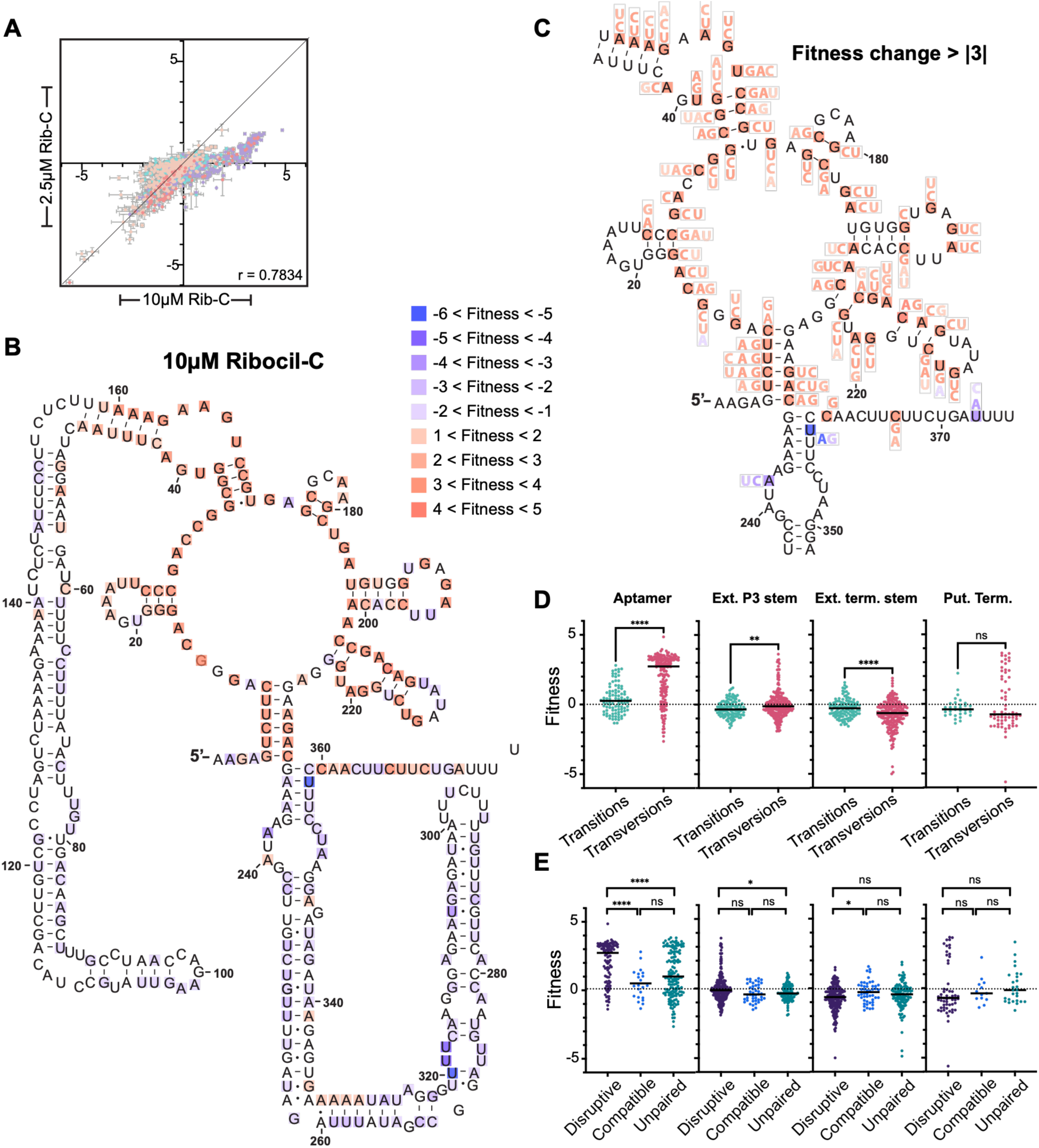
Mutations to the aptamer are beneficial in the presence of ribocil-C. **A)** Fitness values for mutants at 2.5 and 10µM ribocil-C, plotted on x-y axis (error bars correspond to calculated error via DimSum, point color indicates region of the riboswitch corresponding to Fig. 1A). There is a strong correlation between fitness values obtained at the two concentrations (Pearson’s correlation co-efficient r calculated in Graphpad using fitness values in each condition shown on graph, p < 0.0001, Table S4). **B)** Heatmap of mutation fitness in 10µM ribocil-C relative to wildtype mapped to riboswitch secondary structure. Colors displayed at each position correspond to the fitness of the mutation with largest magnitude change from wild-type. Blue: negative fitness, red: positive fitness. **C)** Inset of aptamer and proximal areas highlighting only positions with fitness changes > |3|, all mutations with a fitness change > |1| for each such position are colored corresponding to the fitness change. **D)** Relative fitness of transition versus transversion mutations by region of the riboswitch in the presence of 10 µM ribocil-C shows that transversions confer higher fitness compared to transitions in the aptamer. Populations compared using two-tailed Mann-Whitney test (p>0.05 (ns), p< 0.05 (*), p<0.01 (**), p<0.001 (***), p<0.0001 (****)). Exact p-values for each comparison on Table S3. **E)** Fitness of mutations in the aptamer disruptive of predicted base-pairing are significantly more positive compared to those of compatible mutations or mutations to unpaired bases (see Fig. 1F for predicted secondary structure and Table S2 for classifications of each mutation). Populations compared using a Kruskal-Wallis test with Dunn’s test for multiple comparisons (p>0.05 (ns), p< 0.05 (*), p<0.01 (**), p<0.001 (***), p<0.0001 (****)). Population medians and exact p-values for each comparison on Table S3.

The positively selected mutants at 10µM ribocil-C are enriched in the aptamer compared to the −riboflavin condition **(Fig. 1C, Table S3)**. This difference in fitness is substantial in magnitude (Δmedian = 1.442) indicating that many mutations to the aptamer confer a large fitness advantage in the presence of 10µM ribocil-C. Visualizing the locations of beneficial and detrimental mutations at 10µM ribocil-C on the riboswitch secondary structure shows that a mutation to ∼83% of positions in the aptamer region increases fitness **(Fig. 2B)**, and the majority of mutations conferring large fitness benefits are more disruptive transversions compared to transitions (**Fig. 2C,D, Table S2,S3**). To investigate the role of these changes, we assessed all positions with respect to their predicted base-pairing status. Mutations disruptive to predicted Watson-Crick pairing in the aptamer are more beneficial than mutations compatible with pairing (Δmedian = 2.407), and mutations to unpaired positions that may be in involved with tertiary interactions or ligand binding (Δmedian = 1.900) (**Fig. 2E, Table S3**).

Across other regions of the riboswitch, the relative fitness of mutants has a wider spread compared to the non-selective conditions **(Fig. 1D-F)**. Mutations to the putative terminator, especially transversions disruptive to base-pairing, have a bimodal distribution that includes both strongly positively selected mutants as well as a negatively selected population (**Fig. 2D,E**). The positively selected mutations are typically within the closing stem of the aptamer, and the negatively selected present within the top of the extended terminal stem or the region immediately 5’ of the putative termination site **(Fig. 2B)**. This pattern suggests that mutations that disrupt the aptamer structure or impair its ability to bind ligand are positively selected at 10µM ribocil-C.

### Mutations to the putative terminator region are beneficial at high roseoflavin concentrations

Although both ribocil-C and roseoflavin target the FMN riboswitch and cross-resistance has been observed, they are not similar chemically, and structural studies show different modes of interaction^33^. Furthermore, roseoflavin is thought to be imported via the riboflavin transporter (RibU), under control of this riboswitch^28,34–36^. Given these differences in drug properties, we repeated our experiment with 64nM and 256nM roseoflavin. These concentrations were chosen to be somewhat inhibitory and strongly inhibitory, given the IC50 of wildtype *S. pneumoniae* TIGR4 of 32nM roseoflavin. The cultures showed inhibited growth compared to medium lacking roseoflavin, but ultimately reached the full ∼10 doublings.

At 256nM roseoflavin, most of the reads contained 1-3 mutations, with wildtype only making up ∼10% of the total reads **(Fig. 1B).** The mutations that are strongly positively selected are clustered to the putative terminator region (Δmedian across the region compared to the −riboflavin = 2.067) (**Fig. 1F**, **Fig. 3A,B Table S3**), and many individual mutations display fitness values greater than four **(Table S2**). Furthermore, transversions are significantly more beneficial than transitions (Δmedian = 3.102) **(Fig. 3C, Table S3)**, and specifically mutations disruptive to predicted base-pairing display a positive impact on fitness compared to unpaired nucleotides (Δmedian = 3.128) and to mutations compatible with base-pairing (Δmedian = 3.125) **(Fig. 3E, Table S3)**. Thus, disrupting the base pairing in the putative terminator stem is strongly beneficial in the presence of a high roseoflavin concentration.

**Figure 3:**
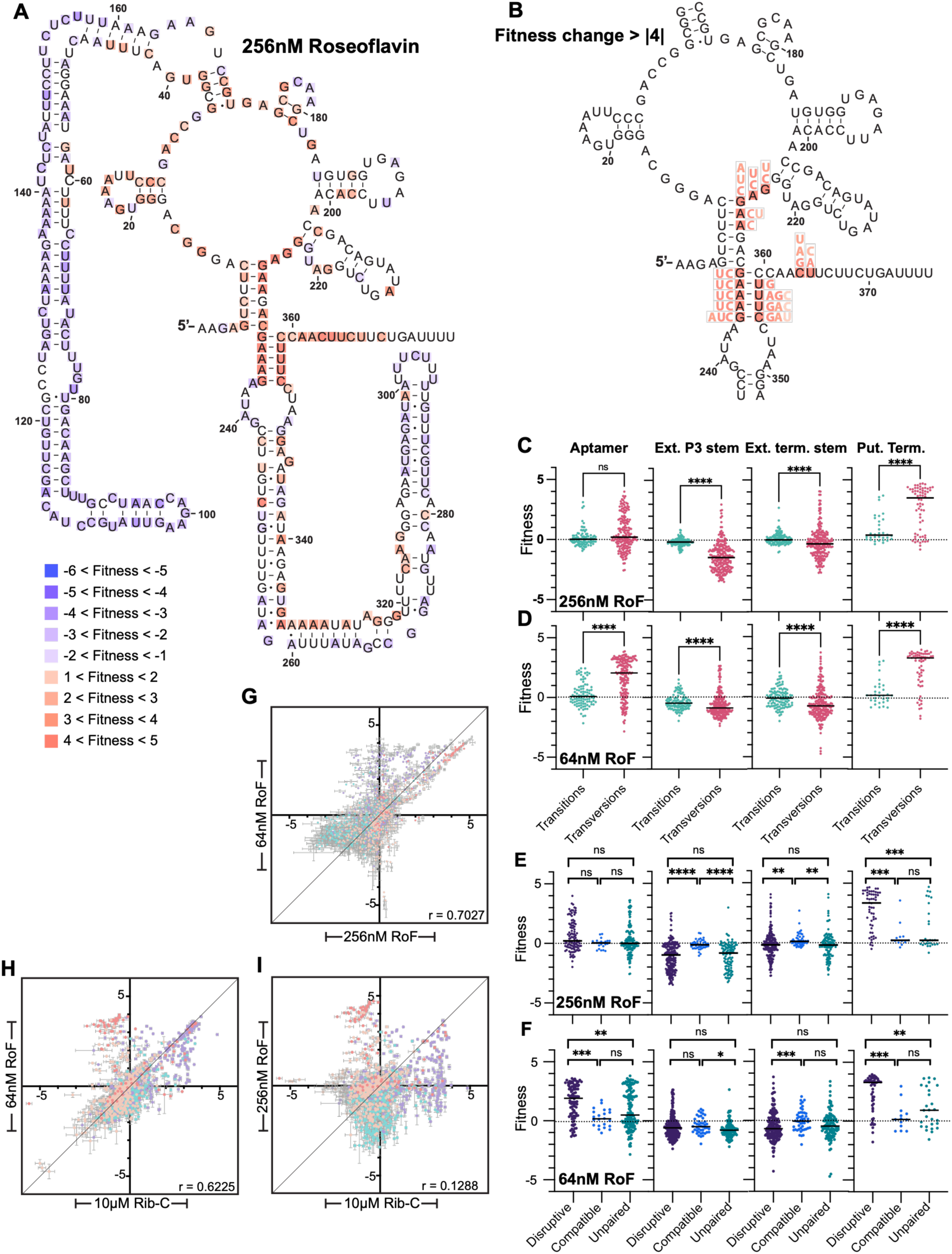
Mutations to the terminator region display the strongest benefit in the presence of roseoflavin, but aptamer mutations may be beneficial at low concentrations. **A)** Heatmap of mutant fitness in 256nM roseoflavin relative to wildtype mapped to riboswitch secondary structure. Colors displayed correspond to the fitness of the mutation with largest magnitude change from wild-type. Blue: negative fitness, red: positive fitness. **B)** Inset of aptamer and proximal areas highlighting only positions with fitness changes > |4|, all mutations with a fitness change > |1| for each position are colored corresponding to their fitness change. All fitness values in Table S2. **C/D)** Fitness of transition versus transversion mutations in 256 nM and 64 nM roseoflavin (statistics as on Fig. 2D) shows a large difference in effects for the putative terminator at both concentrations and for the aptamer only at 64nM. **E/F**) Fitness of mutations by region and base-pairing status in 256nM and 64nM roseoflavin show that mutations disruptive to base-pairing are more disruptive than those that are compatible with the pairing or at unpaired positions in the putative terminator (statistics as in Fig. 2E) **G)** Fitness values from 256nM and 64nM roseoflavin plotted on x-y axis (error bars and point colors as in 2A) show a correlation between the values (Pearson’s r as calculated by Graphpad on graph, p < 0.0001, Table S4). Many aptamer mutations (purple) are poorly correlated. **H)** Fitness values obtained from 64nM roseoflavin and 10µM ribocil-C conditions plotted on an x-y axis as above show that the aptamer mutations positively selected in 10µM ribocil-C correspond with those positively selected in 64nM roseoflavin. **G)** Fitness values obtained from 256nM roseoflavin and 10µM ribocil-C conditions plotted on x-y axis as above show a weaker correlation.

Across the riboswitch there are small magnitude differences compared to our control −riboflavin condition and the spread of relative fitness is larger in the presence 256nM roseoflavin compared to the −riboflavin control condition **(Fig. 1C-E)**. In all regions mutations compatible with predicted base-pairing display a narrower fitness distribution closer to zero compared to mutations that disrupt base-pairing or are predicted to be unpaired (**Fig. 3E**). The highest magnitude fitness changes outside the putative terminator region are to the extended P3 stem **(Fig. 1D)**. These mutations, especially transversions disruptive to predicted secondary structure, are lower fitness while transitions and mutations compatible with base-pairing do not display a substantial change in fitness (**Fig. 3C,E, Table S3)**. This pattern suggests that disrupting the structure of the extended P3 stem reduces *S. pneumoniae* fitness in the presence of roseoflavin.

### Lower roseoflavin concentration reveals a concentration dependent adaptive landscape

In contrast to ribocil-C, the concentration of roseoflavin substantially impacts the relative fitness of mutants. At 64nM roseoflavin the percentage of wildtype reads is slightly higher compared to the 256nM condition (**Fig. 1B**). However, mutations to both the aptamer and putative terminator confer positive fitness values in the presence of 64nM roseoflavin (Δmedians compared to the −riboflavin condition are 1.206 and 2.123 respectively) (**Fig. 1C,F, Table S3**). Comparing the two different antibiotic concentrations, we observe that relative fitness at the two roseoflavin concentrations is correlated (r = 0.7027), but to a lesser degree than the two ribocil-C selected populations (r = 0.7834) (**Fig. 3G, Table S4**). Examination of poorly correlated values reveals that many mutations to the aptamer region display positive fitness at 64nM but are neutral or negative at 256nM roseoflavin. However, mutations to the putative terminator increase fitness at both concentrations.

The aptamer mutations with increased fitness at 64nM roseoflavin are primarily transversions (Δmedian compared to transitions is 1.972) (**Fig. 3D, Table S3**), and the fitness of mutations that disrupt base-pairing is significantly more positive than that of mutations compatible with base-pairing or mutations at unpaired nucleotides (Δmedians = 1.749 and 1.424 respectively) (**Fig. 3F, Table S3**). This pattern suggests that disruptive mutations to the aptamer structure confer a benefit in the presence of lower concentration of roseoflavin, mimicking the pattern observed at 10µM ribocil-C. Comparing the fitness of mutants at 64nM roseoflavin with their fitness in 10µM ribocil-C shows there is significant correlation (r = 0.6225), most notably among mutations to the aptamer (**Fig. 3H, Table S4**). However, a similar comparison of the 256nm roseoflavin condition with 10µM ribocil-C shows few positions with similar behavior in both conditions (r = 0.1298) (**Fig. 3I, Table S4**).

The fitness of mutations outside the aptamer in 64nM roseoflavin is similar to that found in 256nM roseoflavin. Specifically, mutations disruptive to the putative terminator display positive fitness (**Fig. 3D,F Table S3, S4**), with those disruptive of predicted base-pairing displaying higher magnitude values, and mutations to the P3 stem display negative fitness. Thus, the two different concentrations of roseoflavin reveal a pattern where some mutations are beneficial (mutations disruptive to the putative terminator) or detrimental (mutations disruptive to the extended P3 stem) compared to the wild-type sequence at both concentrations, while mutations disruptive to the aptamer are largely beneficial at low concentration, but are more likely to be neutral or negatively selected at a high roseoflavin concentration.

### A biological replicate reinforces findings from technical replicates

To assess the replicability of our findings, we created a second library in the same manner (separate *S. pneumoniae* transformations) as a biological replicate (Lib2). The same amplicon sequencing experiments were repeated for Lib2, and two technical replicates from each culture condition were sequenced (**Table S1, Table S5**). To determine if the biological replicates Lib1 and Lib2 report similar results, we repeated all the analysis and compared the fitness values obtained. For the +riboflavin and −riboflavin conditions, weak or no significant correlation was observed (**Fig. S5C**). All fitness values for Lib1, and very few for Lib2 are > |1| (**Table S2, S5**), thus the poor correlation is most likely due to lack of selective pressure. This absence of selection renders the signal to noise ratio below the point differences in fitness can be used to make definitive conclusions. In contrast, the four antibiotic conditions show significant correlations (p<0.001) with Pearson’s r = 0.5899 to 0.7906 across the four antibiotic perturbed conditions **(Fig. S5, Table S4)**. The lowest degree of correlation occurs in the 2.5 µM ribocil-C condition that is under the least selective pressure. Furthermore, repeating our analyses of transitions/transversions with Lib2 shows similar results to those of our first library (**Fig. S6**) and differences in the statistical significance of comparisons with Δmedian<0.1 across the two replicates bolsters the conclusion that small magnitude differences may not be biologically relevant.

The main difference between the sequenced libraries is the DimSum estimated error on the fitness of Lib2 is greater than that of Lib1 **(Fig. S7, Table S2, Table S5)**. This almost certainly stems from the decrease in technical replicates sequenced from Lib2 compared to Lib1 (**Table S1**), and the reduced read depth that results from this. In addition, in Lib1 in the presence of ribocil-C a handful of mutants in the ext. term. stem confer large fitness defects. However, in Lib2 the same mutants display fitness < |1| (**Fig. S5, Table S2, Table S5**). These findings point toward a trend in our data that the more positive the fitness, the smaller the error, and the more highly correlated the fitness values between biological replicates (**Fig. S5C, Table S2, Table S5**). This is due to the nature of the measurement, where positive fitness values correspond to a higher read count and negative fitness values a lower read count in the raw data. Thus, in our experiments we are more confident of positive fitness values than negative fitness values. Beyond the correlation in fitness values across both libraries, the correspondence in qualitative conclusions drawn from the individual analyses gives us strong confidence in our high-throughput results.

### Positive fitness mutations show divergent behavior in mono- and co-culture

To validate the findings from our high-throughput experiment, we selected a panel of mutations to sample the different regions of the riboswitch found to be beneficial in the presence of high antibiotic concentrations for further analysis. We chose three mutants in different portions of the putative terminator that are strongly beneficial (fitness >4) in the presence of 256nM roseoflavin (RoF1, RoF2, RoF3), four mutants in the aptamer strongly beneficial (fitness >3) in the presence of 10µM ribocil-C (RibC1, RibC2, RibC3, RibC4), and one site with disparate negative and neutral values in the two different replicates (NC1) across all four conditions, and one site neutral to slightly negative in both replicates (NC2) (**Fig. 4A, B)**

**Figure 4:**
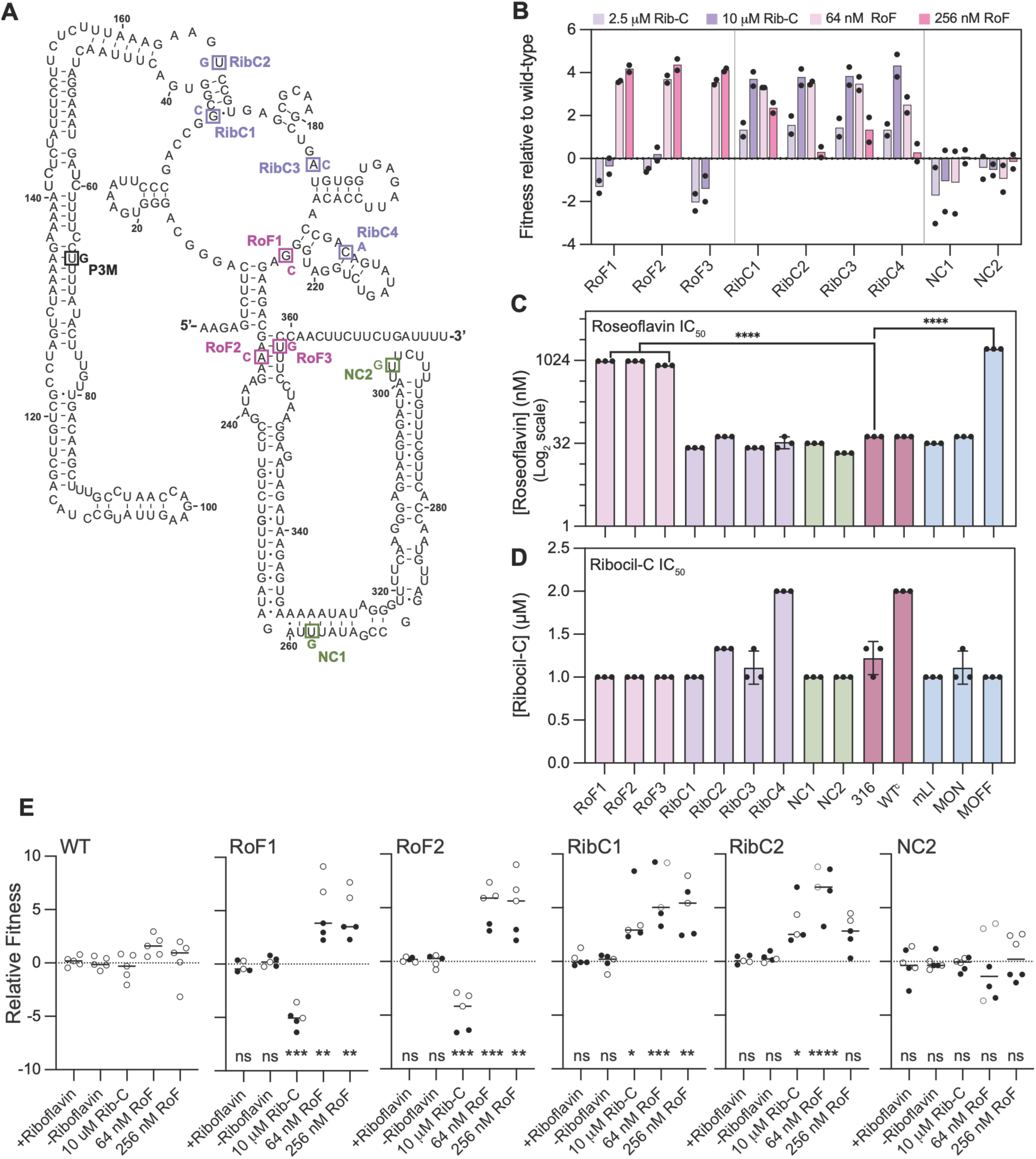
Positively selected mutant phenotypes are replicable via individual competitions with wild-type but are not necessarily antibiotic resistance. **A)** Roseoflavin selected (RoF), Ribocil-C selected (RibC), P3 (P3M), and negative control (NC) mutations shown on the secondary structure of the SP_0488 FMN riboswitch. **B)** Fitness derived from the high-throughput sequencing experiments (Lib1 and Lib2 values indicated by individual points) for the RoF, RibC and NC mutations. **C)** IC_50_ of roseoflavin for RoF, RibC, NC, and additional control strains shows that RoF mutants are resistant to roseoflavin. Pink: RoF selected mutants, purple: RibC selected mutants, green: negative controls, red: parental TIGR4 or WT^C^ control containing only Chl cassette, blue: previously published SP_0488 FMN riboswitch mutants^31^. Each point is a biological replicate (mean of technical triplicate) significance determined via a one-way ANOVA, compared to wildtype 316 (unmodified strain). Only comparisons with significance are indicated (p>0.05 (ns), p< 0.05 (*), p<0.01 (**), p<0.001 (***), p<0.0001 (****)) (See Table S7, growth curves in Fig. S8B,C). **D)** IC_50_ of ribocil-C shows that positively selected strains are not resistant in monoculture. Replicates and data analysis as in part C above. **E)** Co-culture competition assays mirror the results from the high-throughput sequencing experiments. Open points use WT^K^ as the comparison strain, closed points use WT^C^ as the comparison strain. Significant difference in fitness between each strain and the WT^c^/WT^K^ competition for each condition (+/− riboflavin or antibiotic) assessed via one-way ANOVA and Dunnett’s multiple comparisons (p_adj._>0.05 (ns), p_adj._< 0.05 (*), p_adj_<0.01 (**), p_adj_<0.001 (***), p_adj_<0.0001 (****)) (Table S7).

*S. pneumoniae* strains carrying each individual mutation were constructed and validated (see Methods). We assessed five additional strains alongside this panel: the *S. pneumoniae* TIGR4 parental strain (316), and previously published mutants: a strain with a wildtype riboswitch accompanied by a chloramphenicol resistance cassette at the same locus (WT^c^), a ligand-insensitive mutant (mLI), an “always on” mutant (mON), and an “always off” mutant (mOFF)^31^. None of the previously constructed mutants are single point mutations, but rather are larger-scale sequence changes that range from a double mutant (mLI) in the aptamer, to large deletion of the aptamer (mOFF), or putative terminator stem (mON).

The fourteen strains were first assessed for roseoflavin resistance (8nM-16µM). The IC_50_s for the RibC strains, NC strains, mLI and mON are similar to that of the wild-type *S. pneumoniae* (316 or WT^c^). However, the IC_50_s for RoF1, RoF2, RoF3, and mOFF are nearly 30-fold higher at ∼1µM roseoflavin **(Fig. 4C, Fig. S8, Table S7).** Despite their positive fitness at 64nM roseoflavin, the RibC mutants did not display increased resistance to roseoflavin in monoculture. Thus, the mutants to the putative terminator region and the 3’-portion of the aptamer with positive fitness in the presence of roseoflavin are resistant to roseoflavin, but the aptamer mutants, which all display higher relative fitness at the lower roseoflavin concentration, are not resistant.

The same panel of fourteen *S. pneumoniae* strains was assessed for ribocil-C resistance (250nM to 4µM). The IC_50_ for all fourteen strains including the control strains is 1µM – 2µM ribocil-C **(Fig. 4D, Table S7)**. Thus, although the RibC mutants display higher relative fitness compared to wild-type in the context of our high-throughput experiment, they are not resistant to ribocil-C. The benefit conferred by mutations positively selected by ribocil-C is either a more subtle growth benefit that falls short of resistance, or is dependent on the mixed culture of our high-throughput experiment. Despite the lack of change in IC50, we observed that roseoflavin resistant mutants (RoF1, RoF2, RoF3, and mOFF), displayed significantly reduced growth (measured as maximum OD_600_) at 4µM roseoflavin compared to wildtype 316, potentially consistent with the negative fitness values observed in the high-throughput experiment (**Fig. 4B, S8A**).

To address how growth in monoculture vs. co-culture influences the growth patterns of the mutants, we created kanamycin marked *S. pneumoniae* strains carrying the WT riboswitch sequence (WT^k^) and a subset of the positively selected and control mutants (RibC1, RibC2, RoF1, RoF2, NC2) to enable co-culture competition between strains. Using these strains, we conducted co-culture competition experiments for ∼5 doublings in CDM +/− riboflavin, in the presence of 64nM and 256 nM roseoflavin, and 10 µM ribocil-C. The control comparison (WT^k^ vs. WT^C^) shows that the chloramphenicol marked strain has higher fitness in the presence of roseoflavin (median fitness of WT^C^ is 1.2 relative to WT^K^), thus all competitions were conducted in both marker orientations (**Fig. 4E**).

In non-selective conditions (+/− riboflavin), the mutant strains show no competitive advantage compared to the unmutated version (WT). However, the antibiotic conditions do differentiate the strains. RoF1 and RoF2 show a fitness advantage in the presence of both roseoflavin concentrations and negative fitness values in the presence of ribocil-C. RibC1 and RibC2 mutants display positive fitness in the presence of ribocil-C and at 64 nM roseoflavin. RibC2 shows only a non-significant tendency toward higher fitness at 256nM roseoflavin, while RibC1 maintains similar fitness values across both roseoflavin conditions **(Fig. 4E)**, consistent with the higher fitness observed at 256 nM roseoflavin for RibC1 compared to RibC2 (**Fig. 4B**). The NC2 control mutation shows increased fitness variability in the presence of roseoflavin compared to WT, but no significant change in median relative fitness under either antibiotic condition compared to the WT. These findings contrast with antibiotic IC50 measured in culture but mirror the findings of the high-throughput experiment (**Fig. 4B**). Thus, the fitness benefit conferred by our mutations is dependent on a co- or poly-culture environment.

### Mutations beneficial at high roseoflavin and ribocil-C concentrations have opposing effects on gene expression

To connect the phenotypic differences in our strains to gene expression changes, we used ß-galactosidase translational reporter assays to measure gene expression in the presence and absence of riboflavin similarly to previous studies^31^. The wild-type riboswitch represses ß-galactosidase levels approximately 7-fold, consistent with our previous work **(Fig. 5A)**. The RoF mutants, while still displaying riboflavin dependency (2- to 3-fold repression), express the RibU-LacZ translational fusion at much lower (∼10-fold) levels than the riboflavin repressed WT^c^, regardless of whether riboflavin is present in the medium **(Fig. 5A, Table S8)**. In contrast the RibC mutants all constitutively express the RibU-LacZ fusion at higher levels than the riboflavin repressed WT^c^, and often higher than WT^C^ in the absence of riboflavin **(Fig. 5A, Table S8)**.

**Figure 5:**
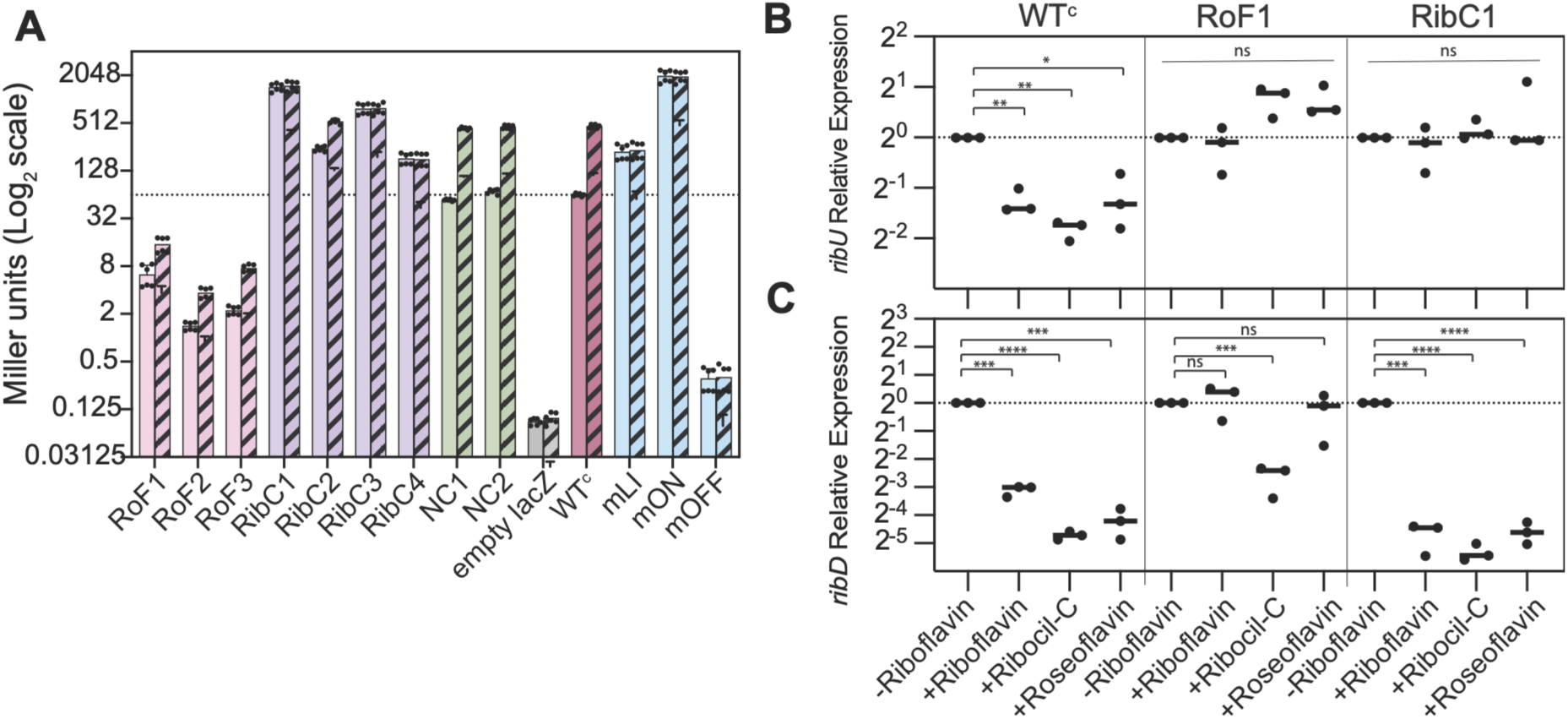
Positively selected mutants display changes in gene expression. **A)** ß-galactosidase assays measure gene expression in the presence (solid bars) and absence (stripped bars) of riboflavin. Error bars represent the standard error of the mean (n=6). Empty l*acZ* (gray) indicates background levels of ß-galactosidase expression in the parental strain. In the presence of riboflavin, RoF mutations display lower gene expression compared to WT (dashed line at WT^C^ + riboflavin) and RibC mutations display higher gene expression. NC mutants do not display behavior statistically distinct from WT^C^, but all other mutants display gene expression in both + and −riboflavin that is statistically different from WT^C^ via a Welch’s ANOVA followed by Dunnett’s test of multiple comparisons (Table S8). **B)** Expression of *ribU* transcript relative to the −riboflavin condition for +riboflavin, +riboflavin, 10µM ribocil-C, and 256nM roseoflavin for WT^C^, RoF1, and RibC1 strains. Points represent biological replicates (n=3) each of which is an average of 3 technical replicates, bar indicates median. Significance assessed via repeated measures one-way ANOVA with Dunnett’s test of multiple comparisons for each strain (Table S8, p_adj._>0.05 (ns), p_adj._< 0.05 (*), p_adj_<0.01 (**), p_adj_<0.001 (***), p_adj_<0.0001 (****)). Expression of *ribU* is mildly (2- to 4-fold) decreased for the WT sequence in the presence of riboflavin, roseoflavin and ribocil-C relative to the −riboflavin condition. Both the RoF1 and RibC1 mutants show no significant change in *ribU* transcript levels to compared to the – riboflavin condition. **C)** Expression of *ribD* transcript relative to the −riboflavin condition for, +riboflavin, 10µM ribocil-C, and 256nM roseoflavin for WT^C^, RoF1, and RibC1 strains (significance as assessed in 5B). For WT^C^ and RibC1, expression of *ribD* is reduced 10- to 40-fold in the presence of riboflavin, roseoflavin and ribocil-C. Expression of *ribD* does not significantly change for the RoF1 mutant in the presence of roseoflavin and riboflavin, and changes more modestly in ribocil-C (∼5-fold repression).

To further examine gene expression, we measured the transcript levels of *ribU* in the WT^C^, RoF1, and RibC1 strains. In the WT^C^ strain, *ribU* levels are modestly repressed (∼2-4 fold) in the presence of riboflavin, roseoflavin, and ribocil-C compared to the absence of riboflavin (**Fig. 5B**). However, both RoF1 and RibC1 strains show transcript levels that are unaltered or slightly increased in response to all three additives relative to the −riboflavin condition **(Fig 5B, Figure S9A),** consistent with our finding that protein expression changes are much lower magnitude in these strains.

To probe how expression of the FMN biosynthetic operon and thus internal FMN concentrations might influence our findings, we also assessed the transcript levels of *ribD* (SP_0178), an FMN biosynthesis gene regulated by the other FMN riboswitch in *S. pneumoniae*. In the wild-type strain we expect this riboswitch to be repressed when the cell has sufficient FMN, or by ribocil-C and roseoflavin, as all three ligands trigger premature termination. As anticipated, *ribD* levels in the wild-type and RibC1 strains are substantially repressed (10- to 20-fold reduction in transcript levels) in the presence of riboflavin, roseoflavin, and ribocil-C compared to the −riboflavin condition (**Fig. 5C, Fig. S9B**). However, the RoF1 strain is not responsive to riboflavin or roseoflavin and *ribD* expression is only ∼5-fold reduced in the presence of ribocil-C compared to −riboflavin. These combined findings suggest that RoF1 expresses RibU at low levels, preventing both riboflavin and roseoflavin from efficiently entering the cell. Thus, FMN biosynthetic genes are not responsive to exogenous FMN, and the strain is resistant to roseoflavin. However, RibC1 expresses the transporter at higher levels, even in the presence of roseoflavin and ribocil-C, and thus constitutively imports riboflavin and roseoflavin, which repress expression of biosynthesis operon.

### Roseoflavin resistant mutants reveal translational repression

While previous 3’-end sequencing data and sequence analysis identified a putative transcription termination site associated with this riboswitch just upstream of the translational start (**Fig. 1A, Fig. S1**)^30^, how this terminator enabled FMN-dependent termination is unclear from the predicted base-pairing. *S. pneumoniae* predominately terminates transcription using intrinsic termination as it does not contain the Rho termination protein^37^, however the site preceding *ribU* does not correspond to a canonical intrinsic terminator despite the presence of a strong stem and polyuridine stretch^38^. Positively selected mutants in the presence of roseoflavin cluster to this region of the RNA, with mutations to positions in the base of the aptamer and the region connecting the aptamer to the ext. term. stem displaying the largest fitness benefits **(Fig. 3B,C Table S2)**. The positively selected mutations are largely expected to decrease the stability of the predicted base-pairing, which in a canonical termination stem typically results in increased read-through to increase gene-expression. However, we find that roseoflavin resistant mutants significantly decrease expression of RibU **(Fig. 5A)**. This unexpected finding motivated us to further assess the entire transcript sequence to determine if there were any potential interactions with sequence following the putative termination site. Investigation of the downstream region revealed the presence of extensive potential base-pairing between the Shine-Dalgarno sequence and the region of the RNA constituting one side of the putative terminator stem and the poly-uridine sequence **(Fig. 6A).**

**Figure 6:**
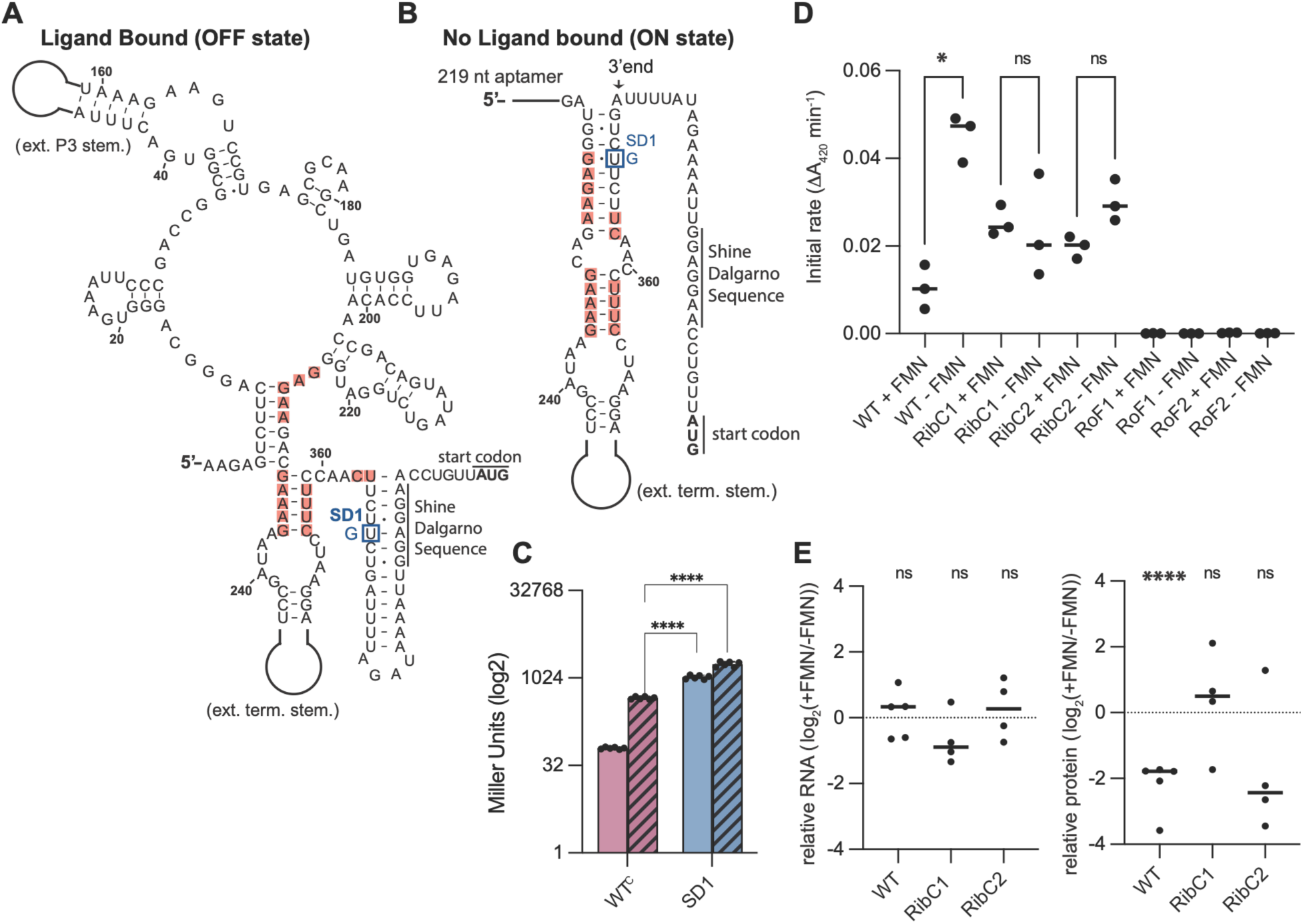
Constitutively repressed roseoflavin resistant mutants suggest a translation inhibition mechanism. **A)** Proposed structure for the ligand-bound form of the aptamer and downstream sequence including the Shine-Dalgarno and start codon indicated. Positions where a mutation results in fitness >4 in 256nM roseoflavin that likely display constitutively repressed expression are indicated. Shine-Dalgarno mutant (SD1) shown in blue. **B)** Proposed structure of the ligand free form of the RNA where the final stem is elongated and the Shine-Dalgarno sequence is released. Positions colored as in A. Positive fitness mutations destabilize this pairing, resulting in increased prevalence of the competing structure shown in A. Shine-Dalgarno mutant (SD1) shown in blue. **C)** ß-galactosidase assays measure gene expression in the presence (solid bars) and absence (stripped bars) of riboflavin. Error bars represent the standard error of the mean (n=6). WT^C^ replotted from Fig. 5A for comparison. In the presence and absence of riboflavin, the SD1 mutant displays a significantly higher gene expression compared to WT^C^ in the absence of riboflavin via Welch’s ANOVA followed by Dunnett’s test of multiple comparisons (p<.0001****, Table S9). **D)** ß-galactosidase activity (initial rate of *o*-nitrophenol production) from *in vitro* transcription/translation reactions shows dependence on FMN for WT sequence (n=3, repeated measures one-way ANOVA with Sidak’s multiple comparisons, Table S9, Fig. S9C). **E)** Ratio in paired +/− FMN reactions of transcript (^32^P-autoradiography) and ß-galactosidase activity (total *o*-nitrophenol production in 30-60 minutes) from the same reaction shows that protein levels are responsive to FMN for the WT sequence, while transcript levels remain more stable (p<.0001****, one sample t-test, theoretical mean = 1.0, n=4 or 5 as shown, Table S9).

The mutations with higher fitness in the presence of roseoflavin destabilize the alternative pairing that frees the Shine-Dalgarno sequence to allow ribosome access **(Fig. 6B)**. This destabilization results in constitutive repression of gene expression as the occluded structure predominates **(Fig. 6A).** Altering bases proximal to the aptamer only destabilize the structure exposing the Shine-Dalgarno sequence (**Fig. 6B)**, while mutations to bases closer to the site of termination (bases 366-372) destabilize both potential structures. Consistent with this, mutations to the putative terminator stem within and immediately following the aptamer (bases 224-236) confer higher fitness than those at the base of the terminator closest to the termination site (bases 354-364). This finding strongly suggests that modulation of translation initiation is a mechanism of action for this riboswitch. To further test the proposed mechanism, we generated a mutant that disrupts the Shine Dalgarno/anti-Shine Dalgaro sequence interaction (termed SD1) in our ß-galactosidase reporter strain (**Fig. 6A, B**). We find that SD1 is constitutively active at higher levels than WT^C^ in both the absence and presence of riboflavin (**Fig. 6C, Table S9**), consistent with this mechanism.

To directly assess whether FMN inhibits translation for this sequence, we performed *in vitro* transcription/translation reactions with the wild-type sequence and several mutants using our ß-galactosidase fusion constructs as DNA template to allow quantitative measurement of protein levels. Initial rates of o-nitrophenyl-β-D-galactopyranoside (ONPG) degradation (o-nitrophenol formation measured as ΔA_420_ min^−1^) from reactions with and without supplemented FMN show that ß-galactosidase levels are reduced by ∼4-fold in the presence FMN for the wild-type sequence **(Fig 6D, S9C)**. For the RibC1 and RibC2 mutant sequences, ß-galactosidase is produced, but the difference between rates in the presence and absence of FMN is not significant. For the RoF1 and RoF2 mutants, there is minimal or no detectable ß-galactosidase activity, consistent with constitutive inhibition of translation.

To assess RNA levels *in vitro*, we performed similar reactions in the presence of trace amounts of α^32^P-GTP and quantified full-length transcript via denaturing PAGE autoradiography. By assessing RNA and protein levels in the same reactions, we find that full-length transcript levels typically vary by <2-fold with the addition of FMN, and endpoint ß-galactosidase assays show a ∼4-fold drop in protein production for the wild-type sequence similar to that observed in the kinetic assays (**Fig. 6E, S9D**). While there are many caveats associated with a non-native RNA polymerase and translation system, these results support the conclusion that FMN-binding inhibits translation for the wild-type sequence, but not mutated sequences that are constitutively expressed. Furthermore, the RoF mutations, conferring reduced protein expression in cells, also display minimal translation *in vitro*, even as full-length transcript is produced (**Fig. S9D**)

### The FMN riboswitch is under weak purifying selection *in vivo*

Using antibiotic perturbation we identified mutations that impact the behavior of the FMN riboswitch. However, these experiments do not ultimately address how such changes impact organismal fitness in a realistic environmental condition. FMN biosynthesis is dispensable for *S. pneumoniae* during *in vivo* infection due to the availability of FMN in the environment. However, the fitness of strains with transposon insertions into *ribU* is attenuated within *in vivo* mouse models^39^. In addition, in our past assessment of fitness conferred by the FMN riboswitch preceding RibU we observed a substantial *in vivo* fitness defect for a truncation mutant that resulted in constitutively repressed expression^31^. To assess whether any single mutant could recapitulate this phenotype, we inoculated mice with our strain library in three different infection groups, allowed the infection to progress for 20-24 hours, and subsequently harvested and homogenized the lung and plated on blood plates to collect the living cells. Genomic DNA was then extracted from colonies collected off these plates and amplicon sequencing performed similarly to our experiment above. A pre-inoculation sample was collected as separate T1 sample for each infection group of mice (T1_mouse_). Close to 70% of the reads coming from both the aggregated T1 _mouse_ and mice infections are the wildtype sequence, similar to the + and −riboflavin conditions **(Fig. 1B)**, suggesting that few mutants are strongly beneficial or detrimental in the *in vivo* infection environment.

When compared to the −riboflavin condition, fitness of mutations to all regions of the riboswitch displayed a larger spread and was slightly negative (**Fig. 1C-F, Table S3**). Transversions are significantly more detrimental to fitness than transitions in all regions with Δmedians ranging from 0.246 to 0.620 **(Fig. 7A, Table S3),** suggesting that more disruptive changes have a more negative impact on fitness across all regions in this environment. Notably, these values are larger than the Δmedians observed under the non-selective + or – riboflavin conditions. The P3 stem shows the largest change in fitness, and mutations disruptive of predicted base-pairing or at unpaired positions are significantly more deleterious than mutations compatible with base-pairing (Δmedians = 0.323 and 0.280 respectively) **(Fig. 7B, Table S3)**.

**Figure 7:**
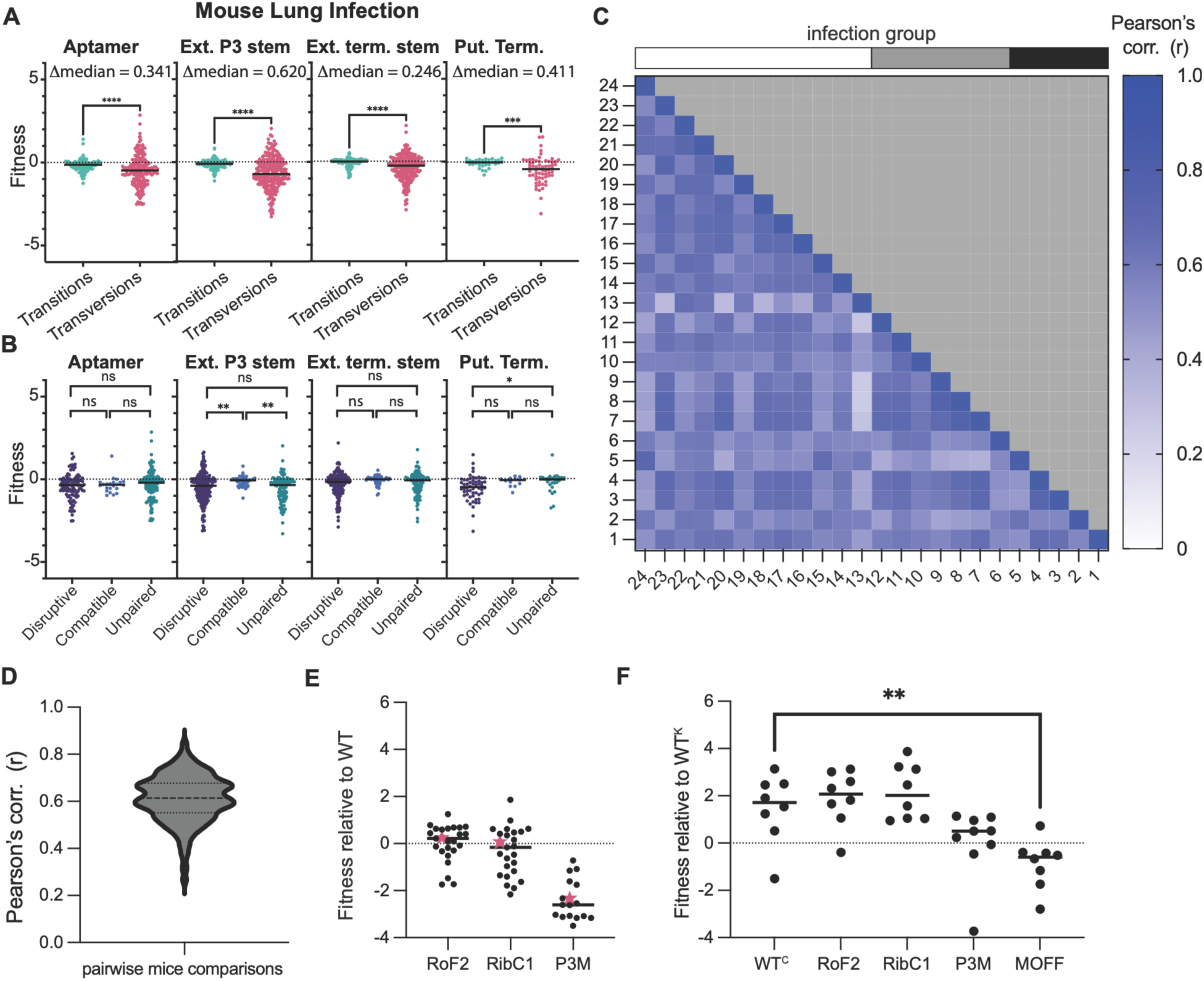
The FMN riboswitch is under weak purifying selection during mouse lung infection. **A)** Fitness of transition versus transversion mutations during mouse lung infection by region of the riboswitch (Aptamer, Extended P3 stem (Ext. P3 stem), Extended Terminal stem (Ext. term stem), and Putative Terminator (Put. Term.)), statistics as in Fig. 2C, Table S3. More disruptive transversions display decreased fitness relative to transitions in all regions of the RNA. **B**) Fitness of mutations in mouse lung infection in the ext. P3 stem that are disruptive to predicted secondary structure or unpaired is significantly lower than that of compatible mutations. Statistics as in Fig. 2D, Table S3. **C)** Correlation of mutant fitness values calculated from individual infected mice (and their paired T1 samples) is reasonably high (median Pearson’s correlation r = 0.61), with no obvious bias exhibited across the three infection groups. **D)** Distribution of Pearson’s correlation coefficient r for all pairwise comparisons in C. **E)** Fitness of selected mutants (RoF2, RibC1, and P3M) within individual mice (black points) and reported fitness from aggregated data (red star). Fewer than 24 points reported for P3M since no counts detected in several mice (Table S2). **F)** Individual competitions between selected strains and kanamycin marked strain WT^K^ within mouse lung infection. Only previously constructed large truncation mutation MOFF^31^ shows a significant difference compared to the WT^C^ control (Kruskal Wallis p = 0.0002, Dunn’s test for multiple comparisons p<.01 ** (Table S7)).

Since positive fitness values tend to be higher confidence than negative fitness values, we sought further corroboration of this finding. The fitness values reported correspond to data aggregated over all mice. To assess the reproducibility and variability of the mouse data, we examined the correlation between fitness values for each mutant determined from the sequencing of individual mice (each compared to the T1 collected alongside its respective infection group). We find that fitness calculated across individual mouse are generally well-correlated, with a median Pearson’s r = 0.61(**Fig. 7C, D**). In addition, there is no obvious clustering to suggest systematic differences between the three infection groups. Thus, while the effects observed for the *in vivo* environment are subtle, they are largely reproducible across separate infections.

To directly assess the reproducibility of observed phenotype, we created a chloramphenicol marked strain carrying a mutation in the P3 stem that displays negative fitness during infection (P3M, **Fig. 5A**, **7E**) and used this strain alongside similar WT^C^, RoF2, and RibC1 strains to conduct co-infection competition assays against the WT^K^ strain that carries no mutations to the riboswitch. As observed in the presence of roseoflavin, the WT^C^ strain displays a fitness benefit compared to the WT^K^ strain during infection (**Fig. 5E**, **Fig. 7F**). Thus, the WT^C^/WT^K^ competition was used as a benchmark for comparison of the mutants. The RoF2 and RibC1 strains display relative fitness similar to that of the WT^C^, mirroring the results of the high-throughput experiment (**Fig. 7E**). The P3M mutation shows a downward trend in fitness that is not statistically significant. We also assessed our previously characterized always OFF mutation, and this strain shows significantly reduced fitness compared to WT^K^, in accordance with our previous findings^31^ (**Fig. 7F**). Thus, we find that single mutations that change gene expression levels and FMN responsiveness have modest or undetectable effects on *in vivo* organismal fitness measured here. Furthermore, the trends observed in the high-throughput experiment are replicable via individual *in vivo* competition assays.

## DISCUSSION

Our experiments have revealed a host of information that informs how RNA function responds to single point mutations in the context of a cellular system as well as uncovered important insights regarding *S. pneumoniae* response to antibiotics and the mechanism of action for this RNA. By monitoring the growth of a strain library of FMN riboswitch variants in seven different environments, we illuminated entire suites of mutations displaying decreased or increased expression and used this data to infer a mechanism of action for the RNA and better understand the complex nature of antibiotic selection and the generation of tolerant variants.

The simultaneous assessment of many mutations allowed us to revise our mechanism of action for this riboswitch. While riboswitch aptamers are typically well-conserved, the features that determine how ligand binding is mediated into genetic regulation or “expression platforms” are not conserved across species and can be challenging or impossible to deduce from sequence analysis alone^40^. The presence of a putative termination site identified via 3’-end sequencing close to the aptamer biased our initial hypotheses toward assuming a purely transcriptional mechanism of action. However, the suite of high fitness mutants in the presence of roseoflavin within the putative terminator lead us to determine that translational repression is an important component of RibU regulation. This finding underscores the challenges associated with prediction of riboswitch mechanism of action in the absence of detailed experiments, even when coupled to high-throughput 3’-end sequencing data.

Beyond specific insights into RNA structure and function, our study also reveals the complex relationship between drugs targeting metabolic control, the native metabolite ligands, and the processes inhibited by such compounds. Constitutively repressed mutants that are strongly beneficial in roseoflavin regardless of concentration (RoF mutants), have a straightforward link to resistance. These mutations reduce transport of both riboflavin and roseoflavin into the cell due to constitutively repressed RibU expression. These strains are thus protected from the negative effects of roseoflavin as the biosynthesis of FMN is not inhibited **(Fig. 4C,5**,**8A)**. This effect mirrors the resistance to roseoflavin displayed by organisms lacking a riboflavin transporter that can be sensitized by exogenous expression of RibU^28^. Our constitutively active mutants (RibC mutants) that result from disruptions to the aptamer confer a relative advantage in the presence of ribocil-C and lower concentrations of roseoflavin. However, these mutants do not confer resistance in monoculture, even as they display a large fitness benefit in both our high-throughput experiment and individual competition experiments (**Fig. 4**).

**Figure 8:**
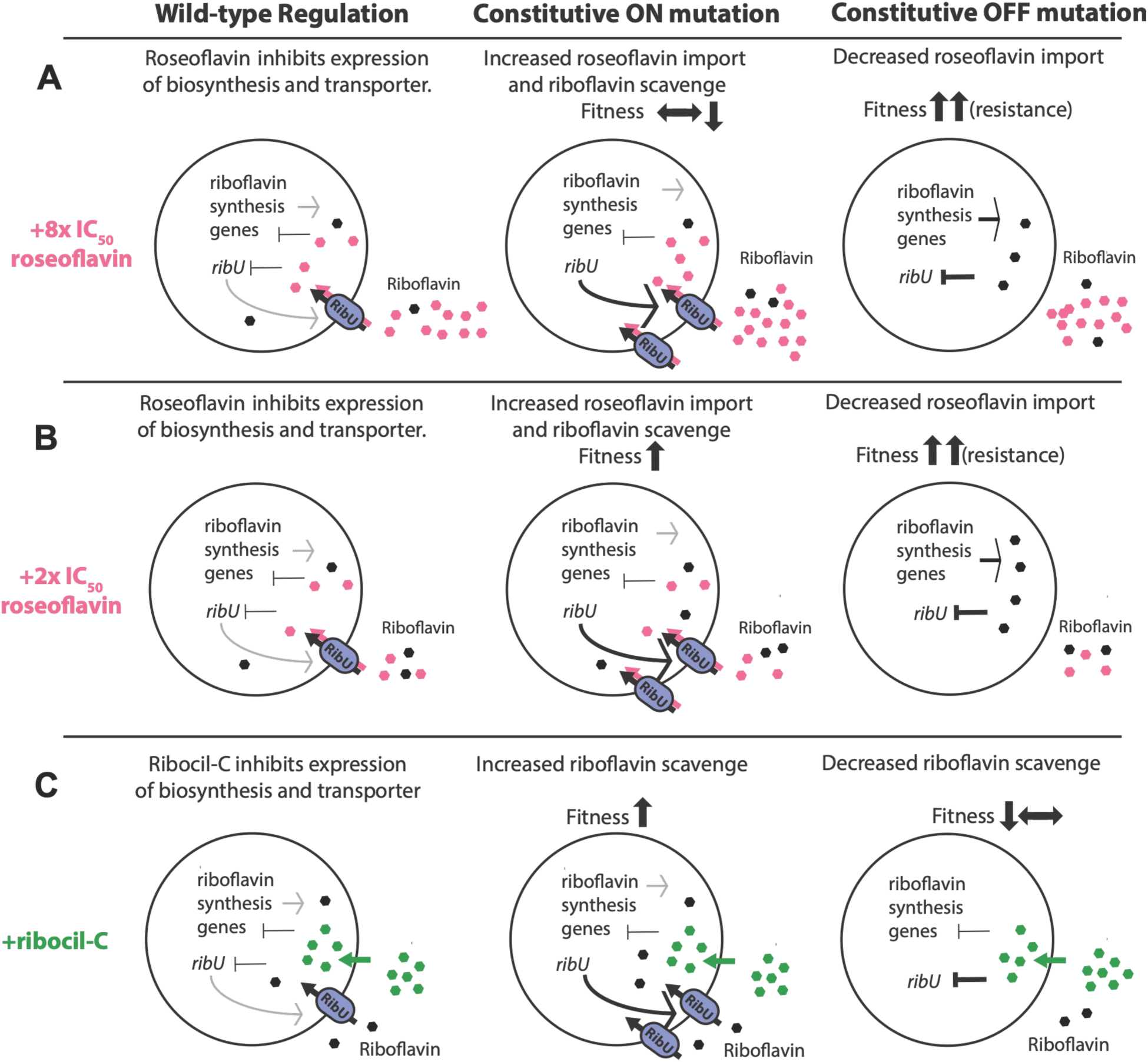
Model for relationship between gene expression and fitness benefits. **A)** For a WT cell in presence of high roseoflavin concentrations (8x IC_50_) synthesis and transport of riboflavin are inhibited. Mutations resulting in constitutively activated RibU expression (ON) have largely neutral or deleterious effects as riboflavin and roseoflavin are both constitutively imported and compete with one another. Mutations resulting in constitutively repressed RibU expression (OFF) substantially reduce import of roseoflavin, resulting in resistance (much higher fitness). **B)** At lower (2x IC_50_) roseoflavin concentrations, mutations resulting in constitutively repressed RibU (OFF) expression confer resistance. Mutations resulting in constitutively activated RibU expression (ON) confer a modest fitness benefit as riboflavin scavenge from the media is increased compared to the wildtype strain, but imported roseoflavin remains relatively low compared to 8x IC_50_ roseoflavin condition. **C)** Ribocil-C enters the cell via a different mechanism. Mutations that result in constitutively activated RibU expression (ON) increase scavenge of riboflavin, increasing fitness, but not enabling monoculture resistance. Conversely, mutations resulting in constitutive RibU repression (OFF) decrease carrying capacity in ribocil-C with neutral to negative effects on fitness.

Based on these combined findings, we speculate that riboflavin scavenge from the medium plays an important role in the fitness of our strains. In an antibiotic treated condition where the WT strain is repressing RibU expression, the constitutively active mutants better scavenge the limited FMN or riboflavin available in the environment, likely derived from autolysis of surrounding cells (**Fig 8B,C**). The ability to continue importing even trace amounts, gives these mutants an advantage in the presence of ribocil-C, which inhibits both FMN biosynthesis and import in the wildtype (**Fig 8C**)^25,26,28^. In this model the increased sensitivity of constitutively repressed mutants (RoF mutants) to ribocil-C reflects their reduced ability to perform such scavenging (**Fig 8C**). This model is also consistent with the fitness advantage of the constitutively active mutants (RibC mutants) at lower roseoflavin concentrations (**Fig. 8B**). At 64nM roseoflavin, we postulate there is a balance achieved where either; (1) constitutive expression of *ribU* allows scavenging in the presence of the drug, or (2) constitutive repression of *ribU* prevents drug entry into the cell and thus biosynthesis is not inhibited. In both situations the concentration of FMN is higher in cells with altered expression compared to wild-type, and thus these cells are positively selected. At higher concentration of roseoflavin, preventing entry of the drug becomes more strongly positive, and enabling scavenge from the medium comes at a larger expense as the drug concentration is higher (**Fig. 8A**).

The most common roseoflavin resistant mutants, across several different bacterial species, map to the aptamer region of FMN riboswitches preceding the biosynthetic operon (*ribD*) ^23,26,28,34,41^. Most mutations are disruptive of RNA structure or ligand binding, result in increased FMN biosynthesis, and when tested display cross-resistance with ribocil-C. However, roseoflavin resistant mutations raised in a *Staphylococcus aureus ΔribD* strain map to the aptamer of the *ribU* FMN riboswitch, but their impact on gene expression was not tested. These mutants display relatively small increases in MIC for both roseoflavin and ribocil-C, and the authors speculate that the modest shifts in susceptibility are due to the role played by the riboflavin:roseoflavin ratio^28^. Our study reinforces that ribocil-C and roseoflavin target both FMN riboswitches. Furthermore, while the most strongly selected mutants at the high concentration of each antibiotic are distinct, there exist a group of mutants that confer a selective advantage in the presence of both antibiotics providing several possible escape routes that extend beyond those typically identified preceding *ribD*.

Our study also shows the extent of natural selection on the riboswitch. We find that single mutations are overwhelmingly neutral in planktonic culture in the absence of targeting antibiotics due to the presence of functional riboflavin biosynthesis machinery. However, within an *in vivo* lung infection model, weak purifying selection is apparent, with many mutations displaying relatively small changes in fitness **(Fig. 7A,B)**. Transversions are always more deleterious than transitions in a pattern that suggests the structure of the RNA plays a role in determining fitness, as transversions (purine to pyrimidine or pyrimidine to purine) are more likely to disrupt Watson-Crick interactions characteristic of RNA structure than transitions (pyrimidine to pyrimidine or purine to purine). However, no single mutation characterized displayed strongly deleterious behavior like that of the previously characterized large deletion mutant^31^.

The biosynthetic operon is arguably masking deleterious changes in transporter expression *in vivo*. However, this attribute showcases the resilience of this pathogen to such perturbations. High-throughput transposon insertion studies show that disruption of riboflavin biosynthesis (*ribB/A,D,E,H*) does not significantly impact fitness, but insertions into *ribU* are deleterious during infection^39^. Furthermore, *ribU* transcript is more abundant than that of *ribD* during infection^31,42^. These data together suggest that transport of riboflavin from the environment is preferred during infection and that biosynthesis is not required for infection fitness. Notably, the truncation mutant (large deletion of entire aptamer and the 5’ portion of the extended stem) displaying significantly reduced fitness also shows reporter expression ∼4-fold lower than that of any RoF mutant (**Fig. 5A**), implying that no single point mutant can inhibit expression this fully. Thus, our findings suggest that this RNA sequence is robust to mutations that are strongly deleterious to the organism.

The strongest selective pressure observed within the *in vivo* studies is in a region that is outside the canonical FMN binding regions of the RNA, the extended P3 stem^21^. Although this sequence is conserved amongst *S. pneumoniae* strains^30^, this feature is not shared in amongst other Streptococcus species suggesting relatively recent introduction^43^. Similar such extensions are common in many structured RNAs, and their origins, the evolutionary pressures upon them, and what if any biological function they may have are not well understood^44^. Here we show this region is subject to purifying selection during infection even as most mutations conferring changes to FMN responsiveness show minimal fitness changes. This combination of results suggests that this insertion may play other roles that remain obscure. There are numerous examples of multiple effectors influencing the same transcript^45,46^, and it may be that the P3 mediates the effect of some other signal, or that its role in tuning gene expression may only be apparent under specific stresses when it modulates the rate of transcription to influence ligand binding or dynamics of RNA structure change.

In conclusion, in this study we start to shed light on the selective pressures necessary to create and maintain complex RNA regulators within biological systems. The conserved complex FMN aptamer is a highly tuned component of an important regulatory pathway in many bacterial species, yet *S. pneumoniae* fitness appears largely robust to single mutation changes to its sequence. However, the assays are conducted over relatively few doublings and thus are likely not sensitive to small changes in fitness that may have large impacts over longer timescales. While our results are confounded by the presence of functional FMN biosynthesis genes, the intrinsic buffering observed reflects the circumstances of a natural pathogen. Thus, the weak purifying selection that we measure is likely to have significant consequences over evolutionary timescales. Under perturbation with two different antibiotics the adaptive landscapes reveal the extent to which the RNA functionality is sensitive to single mutations that result in either constitutive repression or activation of the RNA, as well as illuminate the details of gene-regulation and mechanisms governing antibiotic sensitivity.

## MATERIALS AND METHODS

### Doped oligo library construction

Six doped oligos (1% each non-wildtype base at each position), with 10 bp 5’ and 3’ constant sequences, were ordered from IDT to create a SP_0488 FMN riboswitch mutant library mutated at each nucleotide. For each doped-oligo a DNA second strand was synthesized by annealing the doped oligo with a primer complementary to 3’-end with an overhang in Taq buffer (NEB), followed by addition of dNTPs and Taq polymerase (NEB) and incubation at 72°C for 10 minutes. 3’ and 5’ homology arms were amplified via PCR (Q5, NEB) and purified. The three regions (variable region and two homology arms) for each doped oligo DNA library were then assembled in two Gibson Assemblies: the first with the doped region and the 3’ homology arm, and the second with the 5’ homology arm. The six doped oligo libraries were then mixed together in an equimolar pool and transformed into *S. pneumoniae*^47^ to generate libraries with 10^5^-10^6^ independent colonies. Colonies were scraped from the plates, grown in rich medium (37°C, 5% CO_2_, THY + Oxyrase) for ∼ 2 doublings to OD_600_ ∼1.0. Cells were subsequently gathered by centrifugation and resuspended in 30% volume of fresh THY + 15% glycerol and frozen as individual 250 µL aliquots (starter cultures). Primers used in this study are listed in **Table S10**.

### Amplicon sequencing culture conditions and sample collection

A doped oligo library starter culture was washed three times and used to inoculate 10mL of semi-defined minimal media (SDMM), and grown until mid-log phase. This culture was then washed three times, and used to inoculate three replicates of each experimental condition at an OD_600_ of 0.001. The experimental conditions were in the presence of riboflavin (372 nM), in the absence of riboflavin, 64 nM and 256nM roseoflavin, and 2.5 µM and 10µM ribocil-C, grown in chemically defined media (CDM) without riboflavin^31^. After eight hours of growth (37°C, 5% CO_2_), each culture was spun down, the supernatant pipetted off, and the media replaced with fresh media and ligand. Each culture was grown until it reached an OD_600_ of ∼1, or until 12.5 hours. At this point, the cultures were pelleted, the supernatants removed, and the pellets stored at −80°C unitl genomic DNA preparation.

### *In vivo* FMN riboswitch mutant strain library challenge

Fitness experiments were performed by challenging mice with the FMN riboswitch mutant strain library. Female 6 to 8-week old *M. musculus* (swiss webster) were obtained from Charles River laboratories and allowed to acclimate for 5-7 days prior to infection. Mice were infected in three separate groups (n=8, n=8, and n=16). The strain library was initially grown from a starter culture to mid-log phase, at which point it was washed and resuspended in 1x PBS and two pre-inoculation samples taken for analysis for each group. *M. musculus* were inoculated with 2.5*10^7^ cells via intranasal injection under isoflurane anesthesia, and sacrificed after 24 hours, where the lungs were removed and homogenized in 1x PBS as described previously^31^. 10µl of homogenized lungs were used for serial dilutions to estimate the bacterial burden. The remainder of the homogenized lungs were plated on chloramphenicol blood agar plates, grown overnight, and the resulting colonies collected for genomic DNA extraction and amplicon sequencing. Genomic DNA was collected and analyzed via amplicon sequencing for all animals passing an infection threshold of >1.5×10^5^ cfu recovered (total of 24 animals over the three infection groups).

### Amplicon sequencing preparation

The cell pellets removed from −80°C and gDNA isolated (cell lysis by bead beating followed by Qiagen DNeasy Blood and Tissue kit). DNA quantity was determined using a nanodrop, and each sample was diluted to 50ng/µl. 800ng of gDNA was then used in an initial 400µl 5 cycle PCR, split across 8 PCR tubes, used to add the unique molecular identifiers (UMIs). The products were column purified using the Zymo Clean & Concentrator kit, and the quantity was determined using the nanodrop. 400ng was then used in a 400µl 30-cycle PCR, split across 8 PCR tubes, for amplification of the libraries. These products were then purified using the Zymo Gel Purification kit, and quantified via Qubit. Samples were combined to create an equimolar pool. The pool was quantified using the Qubit, and run on either the NextSeq 2000, using the P1 2×300bp kit, or the MiSeq using the v3 reagent kit. Primers used in this study are listed in **Table S10**.

### Fitness determination

After receiving the Fastq files from the NextSeq 2000 or MiSeq, the paired end reads were initially merged using Vsearch^48^. PCR bias was then corrected for by using AmpUMI twice, the first time using the UMIs attached on the 5’ end, the second time using the UMIs attached on the 3’ end^49^. The deep mutational scanning software DiMSum was then used to identify mutants that are either beneficial, detrimental, or neutral in each condition^32^. DiMSum calculates fitness for each variant compared to the wildtype sequence in each replicate as the natural logarithm of the ratio between sequencing counts in a replicate’s output and input samples relative the wildtype sequence according to equation 1 below and subsequently produces a weighted score across the replicates for each condition.

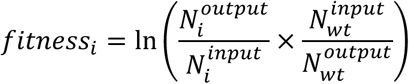

### FMN riboswitch mutant strain construction

Oligos ordered from IDT containing the targeted mutations with additional overhangs for Gibson Assembly were used in an initial 30-cycle PCR to add the mutant and overhang to both halves of the DNA containing the 5’ and 3’ homology arms, the chloramphenicol or kanamycin resistance cassette, and the FMN riboswitch. The PCR products were purified using the Zymo Clean & Concentrator kit, and were used for Gibson Assembly. The Gibson Assembly products were then diluted 1:4, and used in a 30-cycle PCR for amplification. The PCR products were purified using the Zymo Clean & Concentrator kit and quantified using the nanodrop. 50ng DNA was used to transform into wildtype, 316 (TIGR4)^47^. Transformants were plated on chloramphenicol or kanamycin blood plates, and 12 colonies from each mutant were picked, grown for six hours, and analyzed via colony PCR. PCR products amplifying the genome region of interest from promising mutants were sent for Sanger Sequencing (Eton Biosciences), and one from each mutant was selected for use in further experiments. Primers used in this study are listed in **Table S10**.

### Antibiotic Susceptibility Assays

A frozen aliquot of log-phase cells (starter culture) was washed three times in 1xPBS, and then used to inoculate 7.5ml of chemically defined media (CDM) without riboflavin. Cultures were grown for ∼2 hours. The OD_600_ was determined, and cultures were diluted to an OD_600_ of 0.015 in 7ml CDM −riboflavin. Culture and ligand were mixed to obtain the desired ligand concentration (250 nM to 4 uM ribocil-C in 2-fold dilution, 8 nM to 512 nM roseoflavin in 2-fold dilutions, 1 µM to 8 µM roseoflavin in 2-fold dilutions), and 200µl of each condition was plated into three wells of a 96-well plate. The plates were incubated in a BioTek BioSpa 8 (37°C, 5% CO_2_), where their OD_600_ was read every 30 minutes. IC_50_s were determined by identifying the antibiotic concentration at which growth is inhibited >50% compared to growth in the −riboflavin condition at 8.5 hours.

### Individual competitions

For *in vitro* competitions, a starter culture for each strain was washed three times in 1x PBS, and subsequently used to inoculate 5 mL of chemically defined media (CDM) without riboflavin. Cultures were grown for ∼2 hours (OD_600_ ∼0.3), then the WT and a competitor strain with different markers were combined in fresh CDM at OD_600_ = 0.01 of each strain. An aliquot of this mixture was removed and 10-fold dilutions plated for enumeration of the starting ratio. The remaining mixture was diluted 1:1 in CDM +additives to create the following final concentrations of additives: 10 µM ribocil-C, 64 nM roseoflavin, 256 nM roseoflavin, 372 nM riboflavin, and no additive (-riboflavin), and a final starting OD_600_ of 0.01. Cultures were grown at 37°C, 5% CO_2_ for 5-6 hours or until OD_600_ > 0.5. Aliquots removed and 10-fold dilutions plated on blood plates to enumerate relative proportions of kanamycin and chloramphenicol resistant colonies. Fitness calculated from the enumerated colonies as described above.

For *in vivo* mouse competitions, a similar procedure to that described above for *in vivo* library challenge was used. Briefly, each chloramphenicol marked strain (WT^C^, RibC2, RoF1, P3M, MOFF) was combined 1:1 with a kanamycin marked control strain (WT^K^) following initial *in vitro* growth phase and a total of ∼2.5*10^7^ cfu inoculated intranasally into the mouse lung as previously described^30^. Following a 24-hour infection, mice were sacrificed and the lungs removed and homogenized in 1 mL PBS. A series of 1:10 dilutions of the inoculum and homogenate from each mouse lung were plated to enumerate kanamycin and chloramphenicol resistant cfu and fitness calculated as above.

### LacZ reporter strain construction and ß-galactosidase assays

Genomic DNA from strains carrying the RibC, RoF, and NC mutant strains was PCR amplified with primers SP0647+cat_F and lacZ_SP0488_R to generate a product containing the chloramphenicol resistance cassette and the riboswitch through the first three codons of *ribU* under the control of its native promoter. For each mutant these fragments were combined via Gibson Assembly with flanking sequences (amplified by primers SP0647_F/CAT+SP0647_R, and SP0488+lacZ_F/lacZ_R respectively) that correspond to a soluble *lacZ* sequence, and SP_0647, to result in a translational fusion between *ribU* and *lacZ*. These full-length products were PCR amplified and transformed into a previously reported strain wherein the endogenous ß-galactosidase in *S. pneumoniae* TIGR4 (SP_0648) was replaced with a portion of a soluble *lacZ* and a kanamycin resistance marker^31^. Transformants were screened for correct integration of the Chl cassette via PCR and presence of the mutation confirmed via Sanger sequencing of the PCR product. Primers used in this study are listed in **Table S10**. The resulting reporter strains were grown in CDM with 372 nM and 0 nM riboflavin to an OD_600_ of ∼0.5 and harvested. Cells were resuspended in Z buffer (50 mM Na_2_HPO_4_, 40 mM NaH_2_PO_4_, 10 mM KCl, 1 mM MgSO_4_, 50 mM 2-mercaptoethanol) and ß-galactosidase activity assay was performed as described previously^50^.

### qPCR analysis

RNA was isolated from cultures (biological triplicate across different days and medium preparations), with a starting OD_600_ of 0.1, grown for 3 hours in CDM under −riboflavin, +372 nM riboflavin, 256nM roseoflavin, and 10µM ribocil-C conditions, using the Qiagen RNeasy kit (Qiagen). The resulting RNA (200 ng) was treated with DNase (TURBO DNA Free Kit (Ambion)). The DNase treated RNA was used to generate cDNA with Superscript IV Reverse Transcriptase (RT) using random hexamer priming according to the manufacturer’s instructions. Quantitative PCR was performed on 1:2 dilutions in water of both +RT and −RT reactions using Bio-Rad iTaq Universal kit with primers specific to SP_0488 (*ribU*), SP_0178 (*ribD*), SP_2204 (*rplI*), and SP_1219 (*gyrA*) in technical triplicate (**Table S10**). Efficiency of all primer pairs was validated via dilution curves and qPCR data analyzed via ΔΔCT method (**Fig 5BC**) and via 2^−ΔCT^ (**Fig S9A,B**) method to report the relative expression. The geometric mean of both reference genes (SP_2204 and SP_1219) was used for all analyses.

### *In vitro* transcription translation

DNA template corresponding to the full-length FMN riboswitch translationally fused to ß-galactosidase preceded by a T7 promoter was PCR amplified using primers SP0488+T7 and LacZReverse from *S. pneumoniae* reporter strains constructed above (**Table S10**). The DNA product was subsequently purified by a Zymo Clean & Concentrator kit and quantified by nanodrop. For kinetic ß-galactosidase assays 100 ng of this template was used with a PureExpress In Vitro Protein Synthesis Kit (NEB) according to the manufacturer’s instructions (20 µL final reaction volume). This reaction was mixed gently and split into 2 reactions, one of which was supplemented with a final concentration of 2.5 µM FMN. Reactions were incubated at 37°C for 3 hours, and β-galactosidase activity assessed by diluting entire reaction with 90 µL of Z-buffer (see above), adding 10 µL of 4mg/mL ONPG and monitoring o-nitrophenol formation at A_420_ at 30s intervals for 60 minutes in a microtiter plate (**Fig. S9**). Initial rate determined from the linear portion of these curves. For quantification of RNA, similar paired (+/− FMN) reactions were assembled with 1 µL (1:100 dilution) of α-^32^P-GTP (Revvity). Following the 37°C incubation, 4 µL of each reaction was removed and combined with 2x formamide loading buffer (95% formamide, 10 mM EDTA 0.05% bromophenol blue, 0.05% xylene cyanol) for analysis via 4% denaturing PAGE. The gel was dried, exposed to a phosphorimage screen, and ^32^P visualized with a Typhoon FLA 9500. Full-length transcript quantified for each reaction with ImageQuant (Cytiva). β-galactosidase activity from 5 µL of the same reaction was quantified by dilution in 95 µL of Z-buffer, addition of 10 µL of 4 mg/mL ONPG, and incubation at room-temperature for 30-60 minutes until reactions gave visible, but not saturated, yellow color. Reactions stopped with 25 µL 1M NaHCO_3_ at identical elapsed time and A_420_ measured to quantify total activity.

## Data Availability

High throughput sequencing data for this project are available as SRA bioproject: PRJNA1297317. All other data are contained within the supplementary tables.

## Author Contributions

RK: Conceptualization, Methodology, Validation, Formal Analysis, Investigation, Writing – Original Draft, Writing Review & Editing, Visualization. LB: Validation, Investigation, Writing Review. JA: Formal Analysis, Software. EG: Methodology, Software, QF: Data curation, Software. IW: Conceptualization, Methodology, Writing Review &Editing. MM: Conceptualization, Validation, Investigation, Writing Review & Editing, Visualization, Project Administration, Funding Acquisition.

## Animal ethics statement

All experiments conducted in this work were conducted consistently with Boston College IACUC protocol 2022-011 approved on 5/27/2022 and 2025-016 approved on 6/2/2025.

