## Supplemental Figures for "Deep mutational scanning of an *S. pneumoniae* FMN riboswitch reveals diverging adaptive landscapes in response to targeting antibiotics and robustness during infection"

**Figure S1**

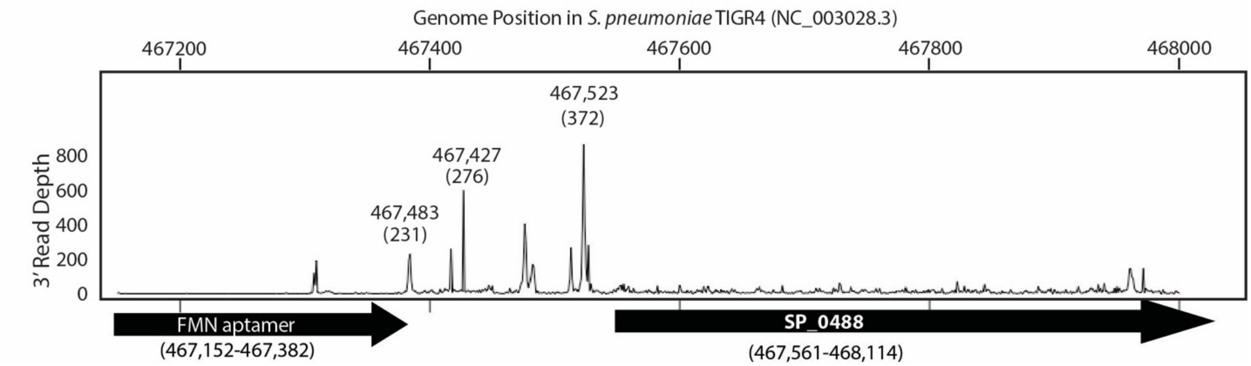


**Figure S1: 3' sequencing data points to a transcriptional mechanism of action for the SP_0488 FMN riboswitch.** Previous 3' sequencing data[30] reveals a potential transcription termination site following the ext. term. stem of the SP_0488 FMN riboswitch in *S. pneumoniae*. Peaks are numbered by both the genome coordinate and according to the aptamer diagram in Fig. 1A.

### Figure S2

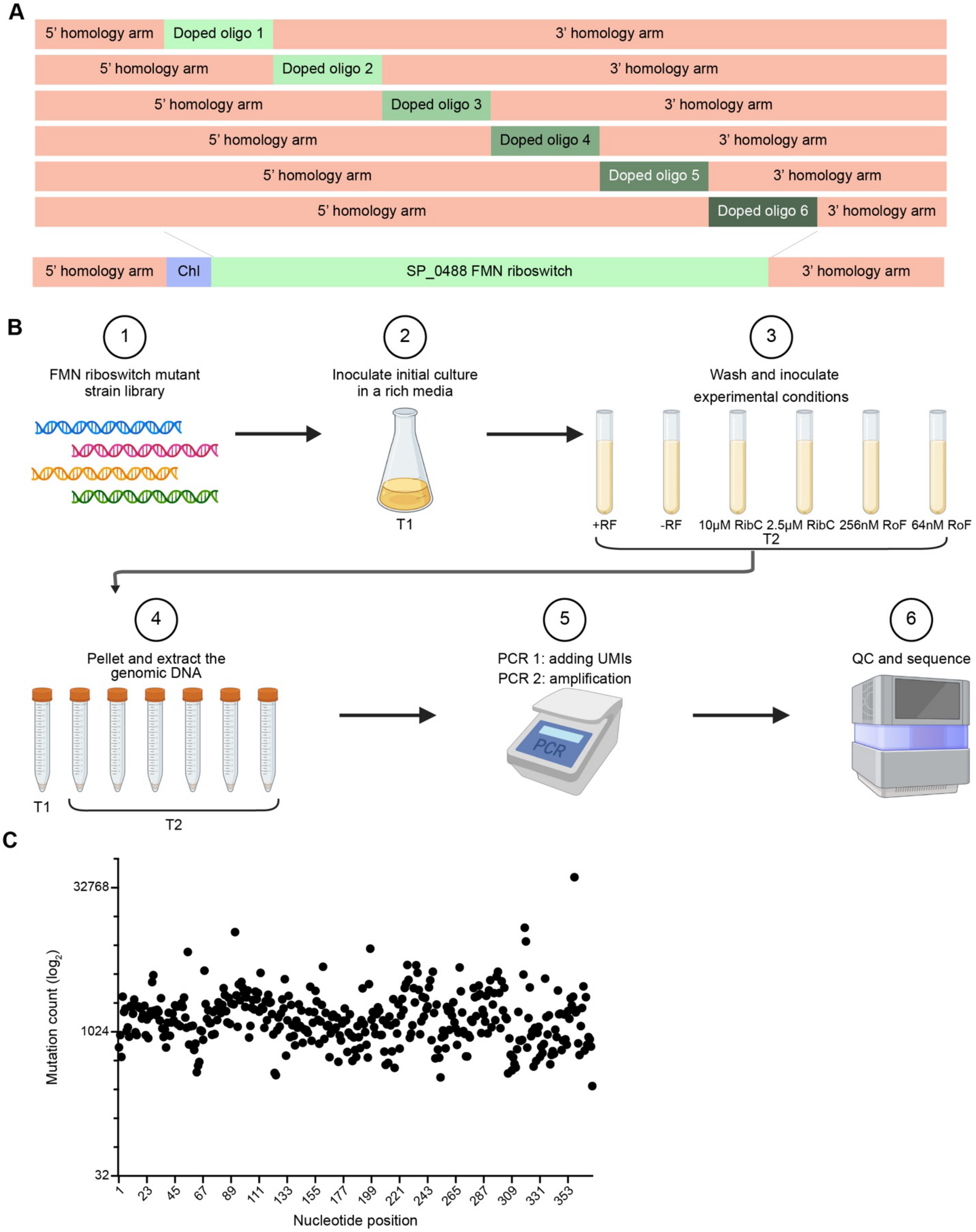


**Figure S2: FMN riboswitch mutant strain library experimental protocol. A)** Six doped oligos with 1% of each alternative base at each position were ordered from IDT and used in second strand synthesis reactions. Gibson Assembly was then used to attach 5' and 3' homology arms that include the chloramphenicol cassette and regions flanking the native locus for the SP_0488 riboswitch. The six libraries were then combined in an equimolar pool, and transformed into *S. pneumoniae* via homologous recombination. **B)** The mutant strain library was inoculated in a rich media and subsequently these cells were washed and used to inoculate six *in vitro* conditions in technical triplicate (three separate cultures grown in parallel from the same medium batch). These cultures grew for ~10 generations, or for 12.5 hours. Cells were collected and used for gDNA extraction. The region of interest was analyzed using two PCRs: the first to add UMIs, and a second for amplification. The amplicon was gel purified, the products were quantified, and sequenced on the NextSeq2000 or MiSeq. **C)** Single nucleotide point-mutation read counts for each nucleotide in the SP_0488 FMN riboswitch at T1 show that each position was mutated in the starting library. Y-axis is represented as a $\log_2$ scale.

**Figure S3**

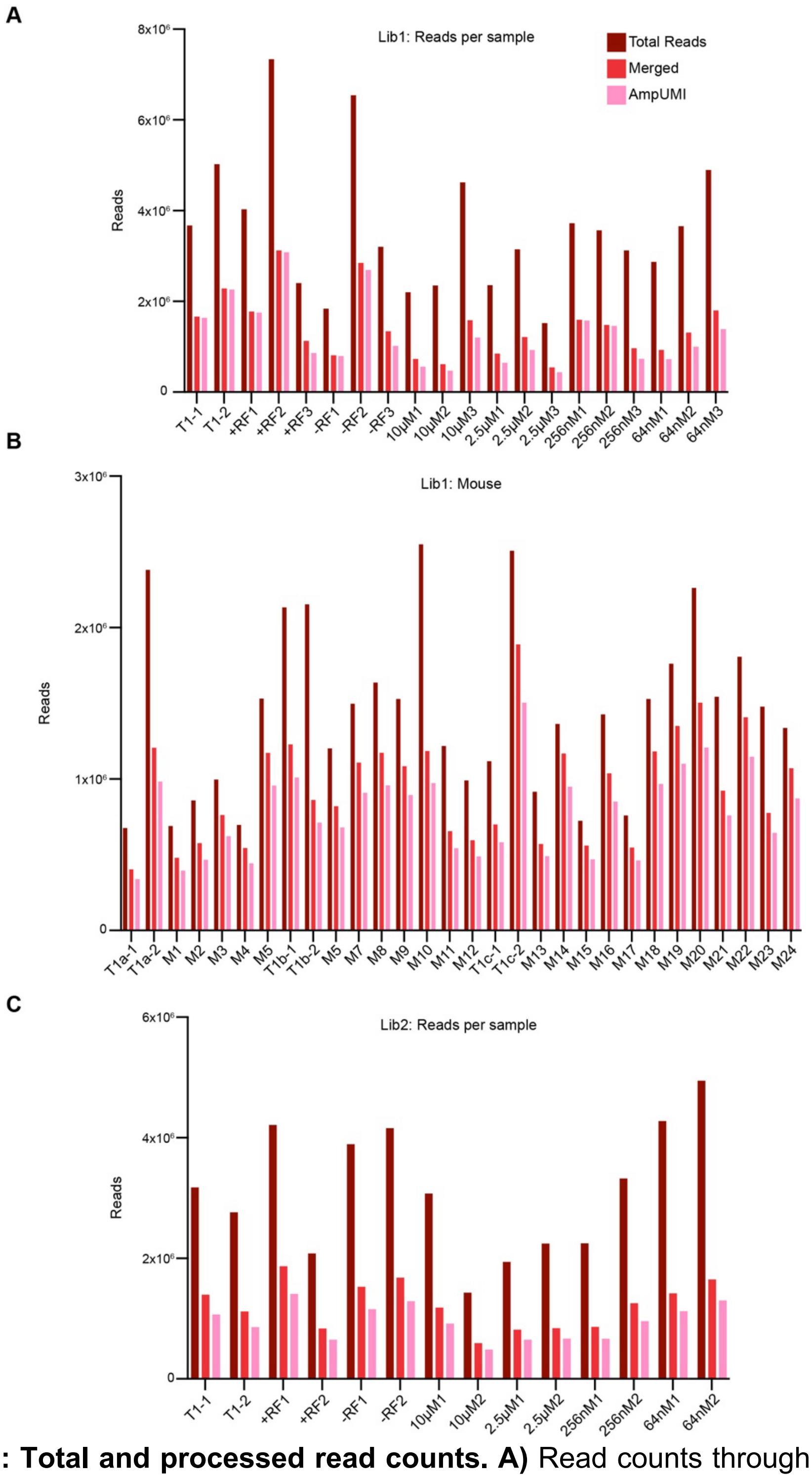


**Figure S3: Total and processed read counts. A)** Read counts through processing for *in vitro* conditions on Lib1 including T1 in duplicate and experimental conditions: +riboflavin, -riboflavin, 10µM ribocil-C, 2.5µM ribocil-C, 256nM roseoflavin, and 64nM roseoflavin in triplicate. **B)** Read counts for each of the 24 mice and the T1 samples collected in duplicate with each set of mice assessed (3 sets total). **C)** Read counts of experimental duplicates on Lib2: T1, +riboflavin, -riboflavin, 10µM ribocil-C, 2.5µM ribocil-C, 256nM roseoflavin, and 64nM roseoflavin. Total reads from the sequencer in dark red, reads after paired-ends are merged in bright red, and reads corrected for PCR bias in pink.

**Figure S4**

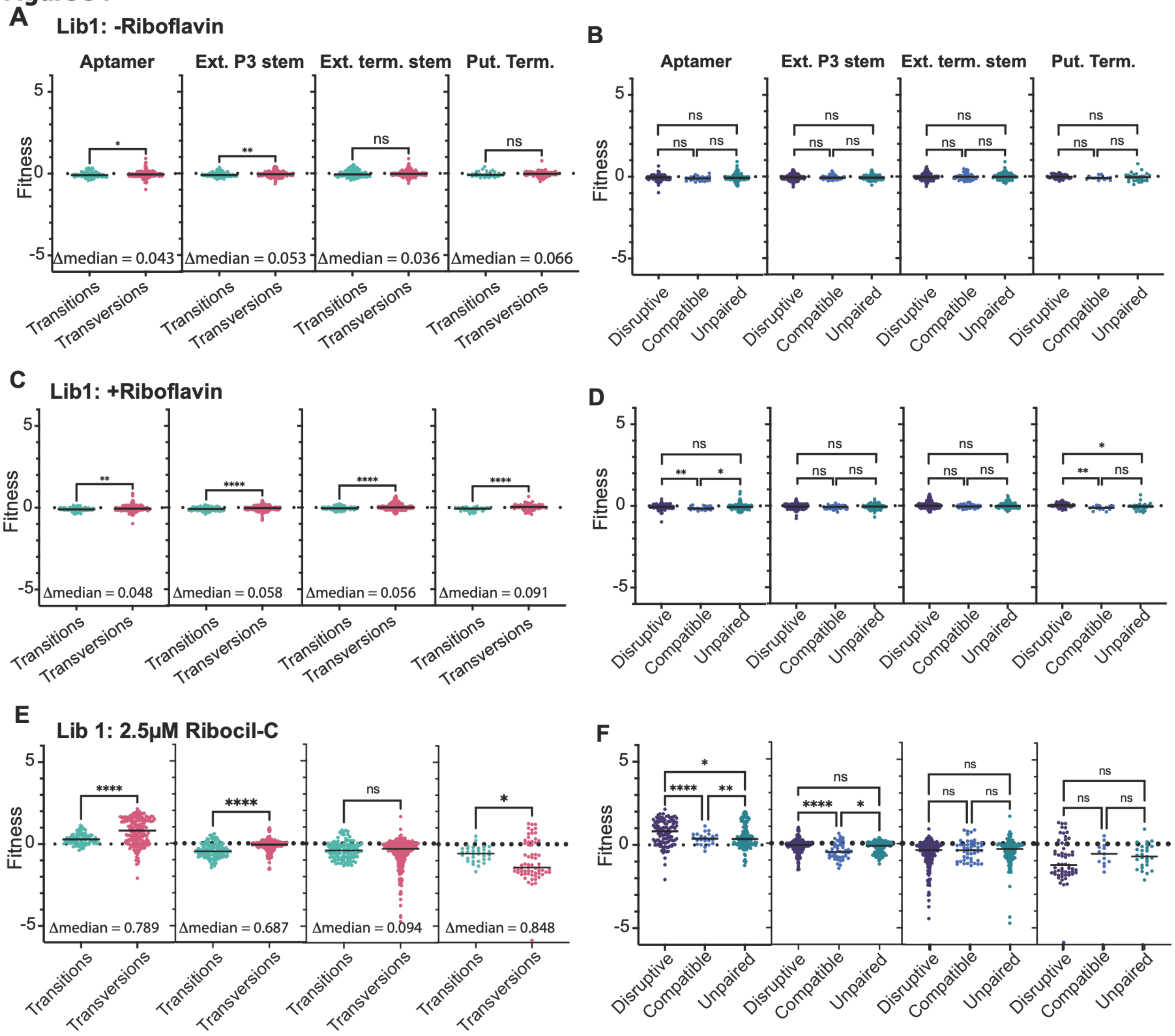


**Figure S4: Most mutations are neutral in the presence and absence of riboflavin. A)** Fitness of transition (purine to purine or pyrimidine to pyrimidine) vs. transversion (pyrimidine to purine or purine to pyrimidine) mutations across different riboswitch regions in the absence of riboflavin. Populations compared using two-tailed Mann-Whitney test (p>0.05 (ns), p< 0.05 (*), p<0.01 (**), p<0.001 (***), p<0.0001 (****)). Exact p-values for each comparison on Table S3. **B)** Fitness of mutations disruptive of predicted base-pairing (see Fig. 1A for predicted secondary structure, Table S2 for classifications), compatible with predicted base-pairing, or not predicted to be Watson-Crick paired (unpaired) in the absence of riboflavin. Populations compared using a Kruskal-Wallis test with Dunn's test for multiple comparisons (p>0.05 (ns), p< 0.05 (*), p<0.01 (**), p<0.001 (***), p<0.0001 (****)). Population medians and exact p-values for each comparison on Table S3. **C)** Fitness of transition vs. transversion mutations across different riboswitch regions in the presence of riboflavin. **D)** Fitness of mutations disruptive of predicted base-pairing compatible with predicted base-pairing, or not predicted to be Watson-Crick paired (unpaired) in the presence of riboflavin. **E)** Fitness of transitions vs. transversions in the presence of 2.5 μM ribocil-C. **F)** Fitness of mutations disruptive of predicted base-pairing compatible with predicted base-pairing, or not predicted to be Watson-Crick paired (unpaired) in the presence of 2.5 μM ribocil-C.

**Figure S5**

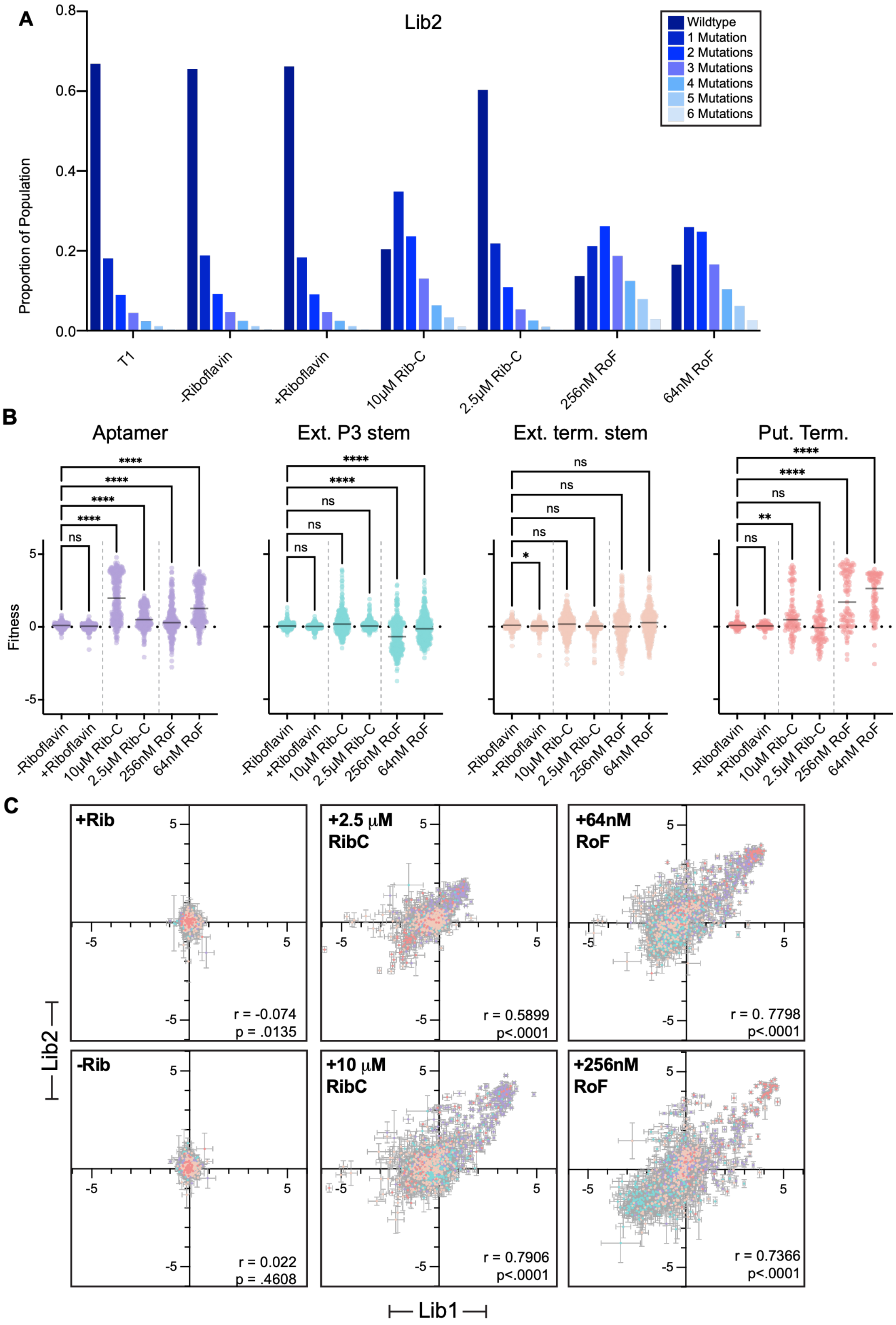
A
Lib2
Proportion of Population
0.0
0.2
0.4
0.6
0.8
Wildtype
1 Mutation
2 Mutations
3 Mutations
4 Mutations
5 Mutations
6 Mutations
T1
-Riboflavin
+Riboflavin
10μM Rib-C
2.5μM Rib-C
256nM RoF
64nM RoF
B
Aptamer
Ext. P3 stem
Ext. term. stem
Put. Term.
Fitness
ns
****
**
*
-Riboflavin
+Riboflavin
10μM Rib-C
2.5μM Rib-C
256nM RoF
64nM RoF
C
+Rib
r = -0.074
p = .0135
+2.5 μM RibC
r = 0.5899
p<.0001
+64nM RoF
r = 0. 7798
p<.0001
-Rib
r = 0.022
p = .4608
+10 μM RibC
r = 0.7906
p<.0001
+256nM RoF
r = 0.7366
p<.0001
Lib2
Lib1

**Figure S5: Lib2 reaffirms data from Lib1. A)** The proportion of processed reads that have 0 (wildtype sequence) to 6 mutations for each condition in Lib2. **B)** Fitness of mutations to different regions of the riboswitch color coded with diagram in Fig. 1A relative to the wildtype sequence in six different environments: chemically defined media (CDM) with no riboflavin (-Riboflavin), CDM+ 375μM riboflavin (+Riboflavin), CDM+ 10μM ribocil-C (10μM Rib-C), CDM+ 2.5μM ribocil-C (2.5μM Rib-C), CDM+ 256nM roseoflavin (256nM RoF), and CDM+ 64nM roseoflavin (64nM RoF). Statistics and analysis as in Fig. 1C-F (Table S6). **C)** Fitness values obtained from two distinct libraries, Lib1 and Lib2, plotted on x-y axis with reported error. There is strong correlation between values obtained for Lib1 and Lib2, with lower error on the positively selected values. Colors correspond to regions of the riboswitch depicted in Fig. 1A.

### Figure S6

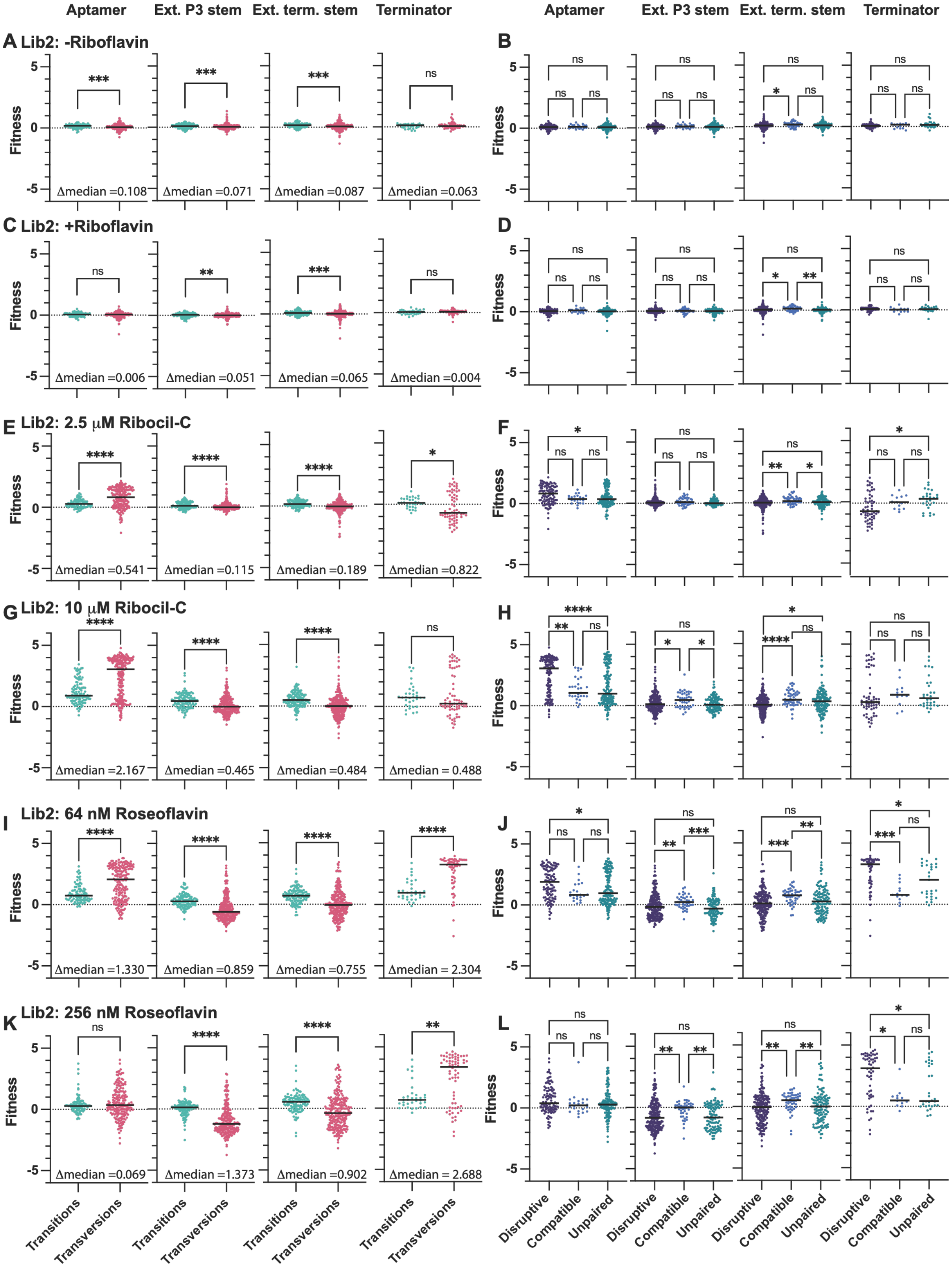

Aptamer
Ext. P3 stem
Ext. term. stem
Terminator
A Lib2: -Riboflavin
B
C Lib2: +Riboflavin
D
E Lib2: 2.5 μM Ribocil-C
F
G Lib2: 10 μM Ribocil-C
H
I Lib2: 64 nM Roseoflavin
J
K Lib2: 256 nM Roseoflavin
L
Fitness
Δmedian =0.108
Δmedian =0.071
Δmedian =0.087
Δmedian =0.063
Δmedian =0.006
Δmedian =0.051
Δmedian =0.065
Δmedian =0.004
Δmedian =0.541
Δmedian =0.115
Δmedian =0.189
Δmedian =0.822
Δmedian =2.167
Δmedian =0.465
Δmedian =0.484
Δmedian = 0.488
Δmedian =1.330
Δmedian =0.859
Δmedian =0.755
Δmedian = 2.304
Δmedian =0.069
Δmedian =1.373
Δmedian =0.902
Δmedian = 2.688
Transitions
Transversions
Disruptive
Compatible
Unpaired

**Figure S6: Lib2 analysis shows similar results that of Lib 1.** Lib2 comparison of transitions vs. transversions (A,C,E,G,I,K) across all conditions and and regions of the riboswitch, analogous to data presented in Fig. 2, 3, and S4 for Lib 1. Lib 2 comparison of mutations that are at unpaired positions, or disruptive or compatible with base-pairing (B,D,F,H,J,L) across all conditions and regions of the riboswitch. Statistics as in Figure 2D,E (Table S6), raw data in Table S5.

**Figure S7**

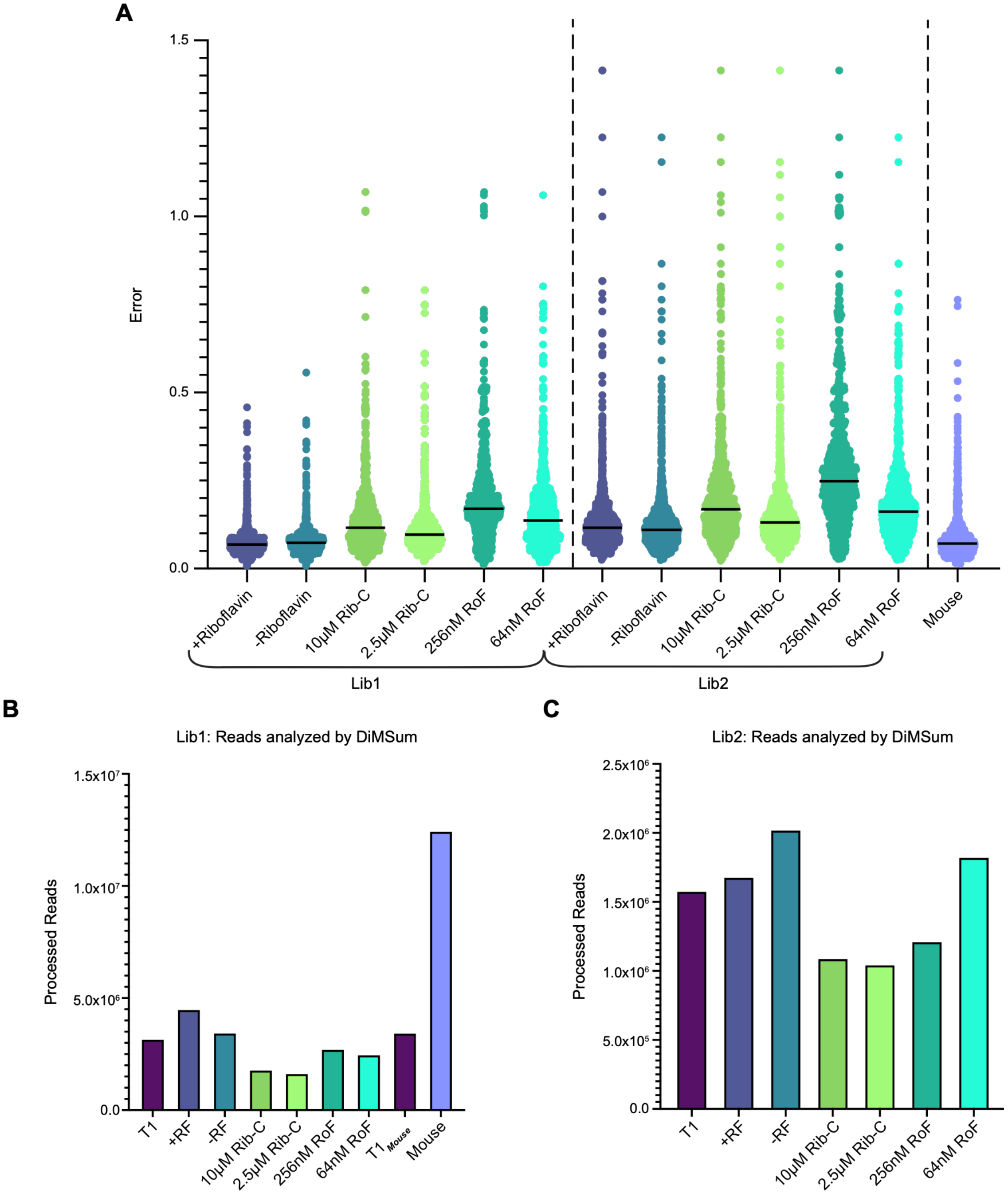


**Figure S7: Lib2 error is greater than Lib1 error when compared by condition. A)** The error on each of the fitness values determined by DiMSum is shown for each condition: -riboflavin, +riboflavin, 10μM ribocil-C, 2.5μM ribocil-C, 256nM roseoflavin, and 64nM roseoflavin, for both Lib1 and Lib2, as well as the error on the sequenced mouse fitness values. **B)** Processed reads that DiMSum was able to analyze from each condition in Lib1, T1, -riboflavin, +riboflavin, 10μM ribocil-C, 2.5μM ribocil-C, 256nM roseoflavin, 64nM roseoflavin, $T1_{mouse}$, and *in vivo* infection. **C)** Processed reads that DiMSum was able to analyze from each condition in Lib2, T1, -riboflavin, +riboflavin, 10μM ribocil-C, 2.5μM ribocil-C, 256nM roseoflavin, and 64nM roseoflavin.

### Figure S8

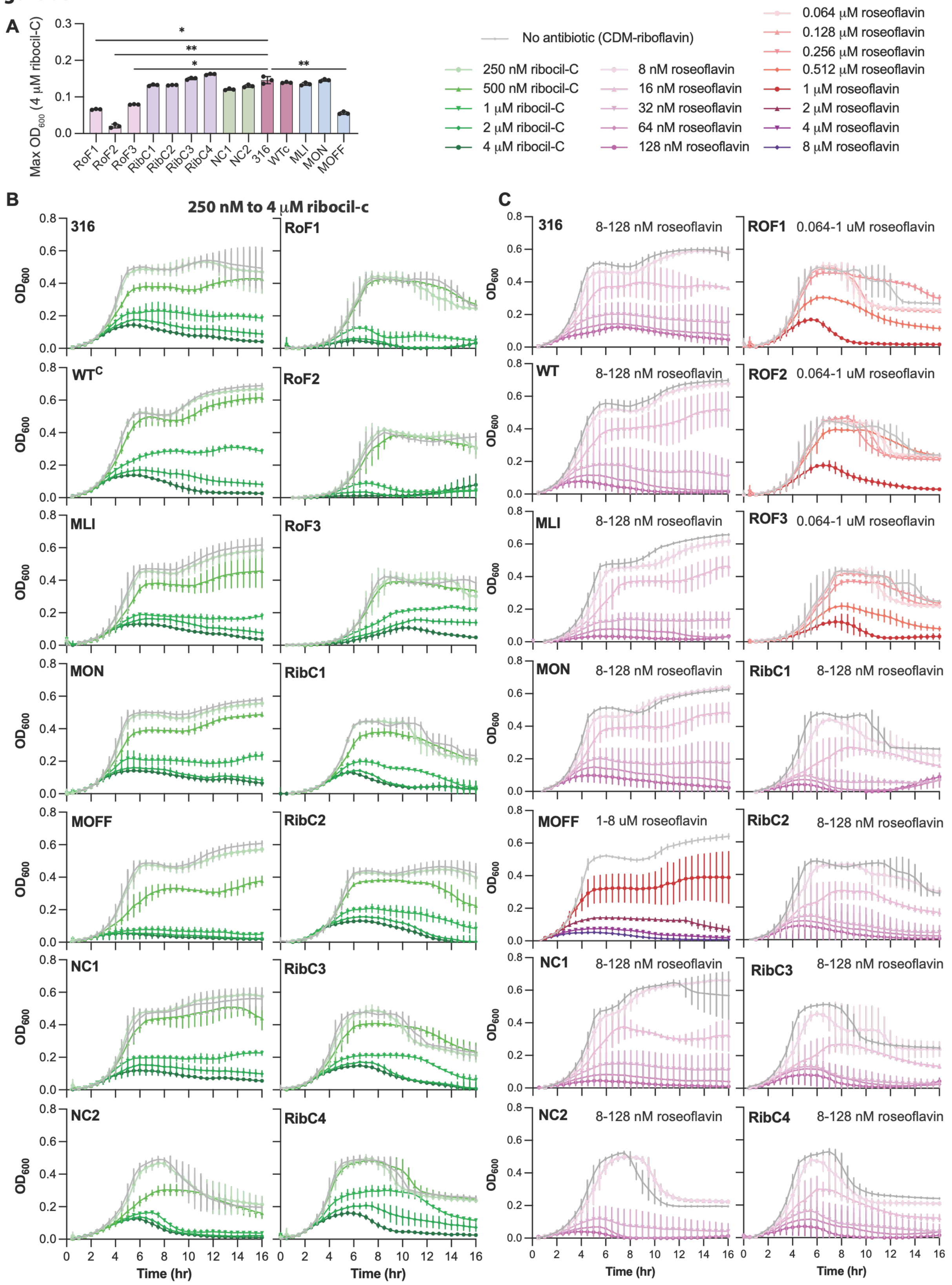

A
Max OD600 (4 μM ribocil-C)
RoF1
RoF2
RoF3
RibC1
RibC2
RibC3
RibC4
NC1
NC2
316
WTc
MLI
MON
MOFF
No antibiotic (CDM-riboflavin)
250 nM ribocil-C
500 nM ribocil-C
1 μM ribocil-C
2 μM ribocil-C
4 μM ribocil-C
8 nM roseoflavin
16 nM roseoflavin
32 nM roseoflavin
64 nM roseoflavin
128 nM roseoflavin
0.064 μM roseoflavin
0.128 μM roseoflavin
0.256 μM roseoflavin
0.512 μM roseoflavin
1 μM roseoflavin
2 μM roseoflavin
4 μM roseoflavin
8 μM roseoflavin
B
250 nM to 4 μM ribocil-c
C
8-128 nM roseoflavin
0.064-1 uM roseoflavin
1-8 uM roseoflavin
ROF1
ROF2
ROF3
WT
OD600
Time (hr)

**Figure S8: A)** Maximum $OD_{600}$ for strains when grown with 4μM ribocil-C for 8.5 hours shows that RoF mutants, and mOFF are more susceptible than the unmodified strain to ribocil-C. Each point represents mean of technical triplicates. Data analyzed via a nested one-way ANOVA, compared to wildtype 316 (unmodified strain). All strains without significance indicated are not significant (ns) when compared to wildtype 316, ($p>0.05$ (ns), $p< 0.05$ (*), $p<0.01$ (**), $p<0.001$ (***), $p<0.0001$ (****)) (See Table S7). **B)** Growth curves used to calculate ribocil-C IC50 (Fig. 4D). Shown is the average and standard error of three biological replicates (each collected in technical triplicate). **C)** Growth curves used to calculate roseoflavin IC50 (Fig. 4C). Shown is the average and standard error of three biological replicates (each collected in technical triplicate).

Figure S9

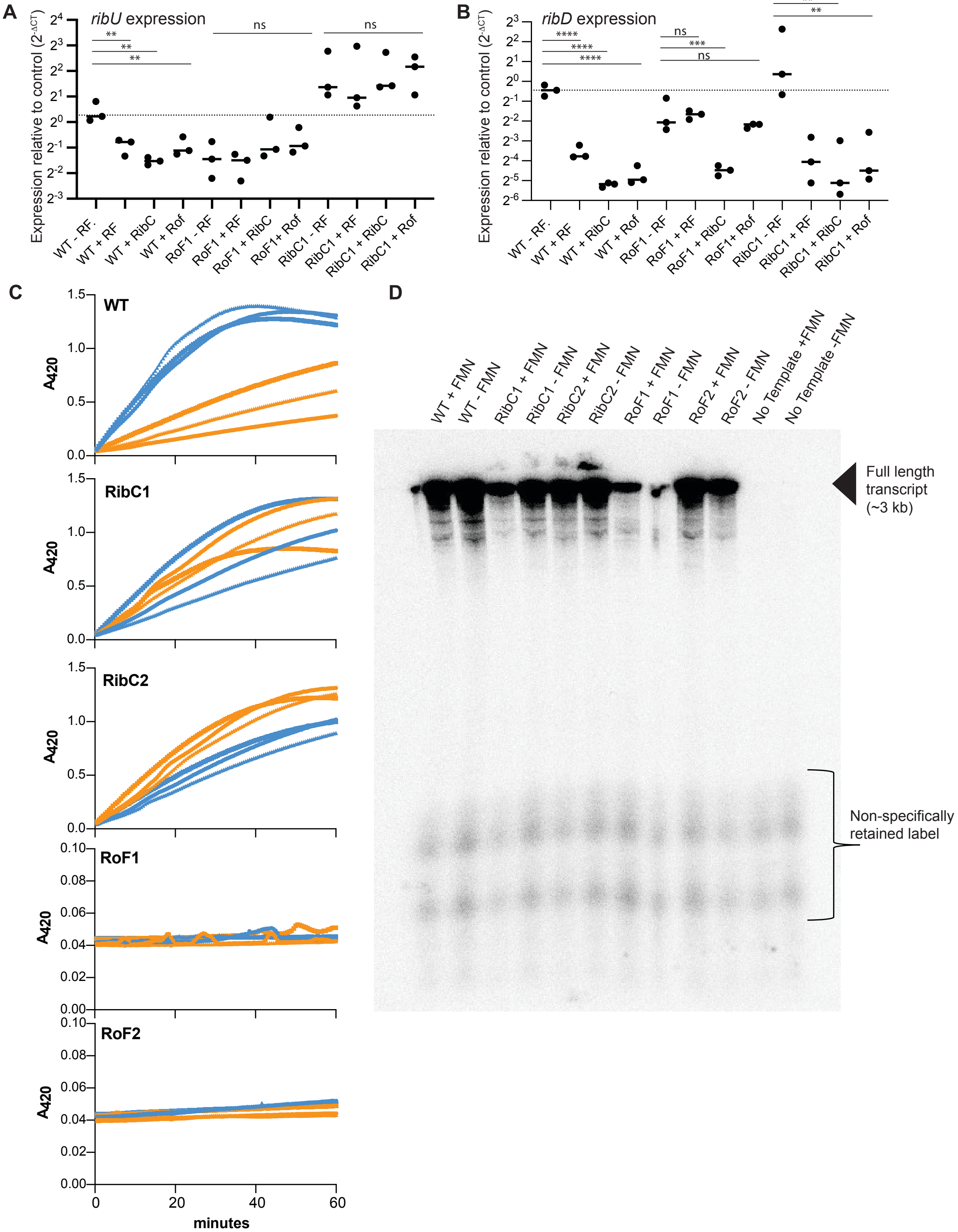


**Figure S9: A)** *ribU* transcript levels compared to control transcripts calculated via $2^{-\Delta CT}$ method. Dashed line indicates WT expression in the absence of riboflavin. **B)** *ribD* transcript levels compared to control calculated via $2^{-\Delta CT}$ method. Dashed line indicates WT expression in the absence of riboflavin Significance determined via one-way ANOVA and Dunnett's test for multiple comparisons (* $p<.05$, ** $p<.01$, *** $p<.001$, **** $p<.0001$, Table S8) **C)** ß-galactosidase activity for WT, RibC1, RibC2, RoF1, RoF2 measured via observation of ONPG degradation to o-nitrophenol at $A_{420}$ at 30s intervals for 60 minutes. **D)** Example autoradiogram of denaturing 4% PAGE gel showing $^{32}P$ incorporation into full-length transcript for all mutants, but no product for no template controls.
